# Condensates in RNA Repeat Sequences are Heterogeneously Organized and Exhibit Reptation-like Dynamics

**DOI:** 10.1101/2021.02.20.432119

**Authors:** Hung T. Nguyen, Naoto Hori, D. Thirumalai

## Abstract

Although it is known that RNA undergoes liquid–liquid phase separation (LLPS), the interplay between the molecular driving forces and the emergent features of the condensates, such as their morphologies and dynamical properties, is not well understood. We introduce a coarse-grained model to simulate phase separation of trinucleotide repeat RNAs, which are implicated in neurological disorders such as Huntington disease and amyotrophic lateral sclerosis. After establishing that the simulations reproduce key experimental findings (length and concentration dependence of the phase transition in (CAG)_n_ repeats), we show that once recruited inside the liquid droplets, the monomers transition from hairpin-like structures to extended states. Interactions between the monomers in the condensates result in the formation of an intricate and dense intermolecular network, which severely restrains the fluctuations and mobilities of the RNAs inside large droplets. In the largest densely packed high viscosity droplets, the mobility of RNA chains is best characterized by reptation, reminiscent of the dynamics in polymer melts. Our work provides a microscopic framework for understanding liquid–liquid phase separation in RNA, which is not easily discernible in current experiments.

## Introduction

The discovery that germline P granules form liquid droplets resembling compartments without membranes^1^ has resulted in a revolution, providing an impetus for a deeper understanding of how the cytoplasm and the nucleus are organized. In turn, this finding has inspired a large number of studies in a variety of unrelated systems that undergo liquid–liquid phase separation (LLPS), postulated to be the mechanism by which such compartments form.^2–14^ A vast majority of such studies have focused on liquid organelles composed of intrinsically disordered proteins/regions (IDP/IDR) or IDPs interacting with RNA molecules that could engage in multivalent interactions.^5^ These findings are typically explained using analogies to sol-gel phase transitions known in synthetic polymers, although there are differences due to the heteropolymer nature of biological sequences.^4, 15–18^ However, the molecular details of how proteins and nucleic acids, with diverse sequences, coalesce and undergo LLPS to form condensates, and potentially mature into gel-like or even ordered states, are still elusive. Furthermore, the conformations and dynamics of the molecules inside the condensates are essentially unknown either in experiments or in simulations.

That interactions between RNA and IDP, such as FUS containing RNA binding C-terminal domain and low complexity N-terminal IDP domain, promote phase separation is well documented.^19–21^ It is also known that RNA plays an important role in driving the formation, stability and morphology of the biomolecular condensates.^6, 12, 13, 21–31^ Recent experiments have also shown that RNA alone is sufficient to self-organize into condensates and gel-like states *in vitro*.^27, 32, 33^ For example, poly-rU with a small addition of short cationic polypeptides can drive reversible LLPS.^27, 32, 34^ Even in germ granules, containing proteins, there is evidence for well organized RNA clusters stabilized by homotypic interactions.^35, 36^ Despite the importance of RNA–RNA interactions in driving LLPS, not much is known quantitatively about the organization and dynamics of RNA condensates.

Nucleotide repeat expansion has been known to cause several neurological and neuromuscular disorders such as Huntington disease, muscular dystrophy and amyotrophic lateral sclerosis.^37–40^ The disease-associated repeat sequences have been found in both the coding and non-coding regions of the transcripts.^38^ However, determining exactly how these repeat expansion sequences cause diseases is still elusive. Recently, Jain and Vale (JV)^33^ showed that the trinucleotide repeat sequences (CAG)_n_, (CUG)_n_ and repeats of the hexanucleotide (G_4_C_2_) form biomolecular condensates. The main findings of the JV study are: The high GC content sequences (CAG)_n_, (CUG)_n_ as well as G_4_C_2_ repeats form droplets, which over time, transform to a gel-like state at high RNA concentrations. (ii) Phase separation occurs only if the number of repeats exceeds a critical value, which is similar to the value where diseases appear.^33, 41^ (iii) *In vivo*, the gel-like state is abolished (except in the repeats of G_4_C_2_), and only a liquid-like state is found. Presumably, this is due to active forces generated by an energy source (ATP binding or hydrolysis, for example). Although RNA transcripts are usually exported to the cytoplasm, the (CAG)_n_ foci, once formed, are preferentially retained in the nucleus and co-localizes with nuclear speckles that sequester splicing factors. These findings provide the needed impetus for investigating the relationship between the tendency to undergo LLPS and the neurotoxicity of the repeat RNA sequences.

In order to provide insights into the mechanism of condensate formation of the RNA repeat sequences, the organization of the RNA chains in the condensates and the associated dynamics, we created a minimal coarse-grained model, representing each nucleotide by a a Single Interaction Site (SIS), to investigate the molecular mechanism of LLPS in RNA repeat sequences. We incorporated only features that are essential for intra- and intermolecular interactions of RNA. In creating the model, we adopted a top-down approach with the goal that it be sufficiently simple so that multi-chain simulations are feasible. The resulting SIS model has only one parameter that sets the energy scale for base pair interactions, which was chosen to reproduce the known structures of a short (CAG)_2_ duplex.^42^ We then performed multi-chain simulations to decipher the LLPS mechanism in the sequence studied by JV *without adjusting any parameter* to match experiments. We quantify the concentrations of the two phases and show unambiguously that phase separation indeed occurs in the simulations, which accords well with experiments. Our simulations recapitulate the length and concentration dependence of the phase separation, in agreement with the *in vitro* experiments. We show that intermolecular base pair interactions drive phase separation. Unexpectedly, we find that once RNA molecules are recruited in the droplets, they undergo large conformational change, from a hairpin-like conformation in isolation to a stretched state in order to form an extensive network of intermolecular interactions. This soft network, in turn, constrains the RNA conformational fluctuation and mobility. The RNA chains in the high density viscous droplets are conformationally and dynamically heterogeneous. In the largest densely packed high viscosity condensates the RNA chains move predominantly by “slithering” along their contour lengths. Such motions are reminiscent of reptation in polymer melts envisioned 50 years ago by de Gennes.^43^ Our work provides important microscopic details of the mechanism of LLPS in the RNA repeat sequences, which currently are not easily accessible in experiments.

## Results

### LLPS depends on RNA concentration and size

Fig. 1**a** illustrates the transition from monomers to condensates as a function of time, *τ*, for the (CAG)_47_ polymer. At *τ* = 0, the CAG chains exist as monomers. At intermediate times, oligomers form, which subsequently fuse together resulting in large droplets as time progresses (see also Movie S1 for a vivid illustration of the phase separation process). The fraction of RNA chains in the oligomers increases, reaching a peak value at intermediate times, and diminishes at long times. The increase in the droplet size coincides with a decrease in the number of RNA chains in the oligomers, suggesting that the small clusters must fuse to form large droplets.

**Figure 1.**
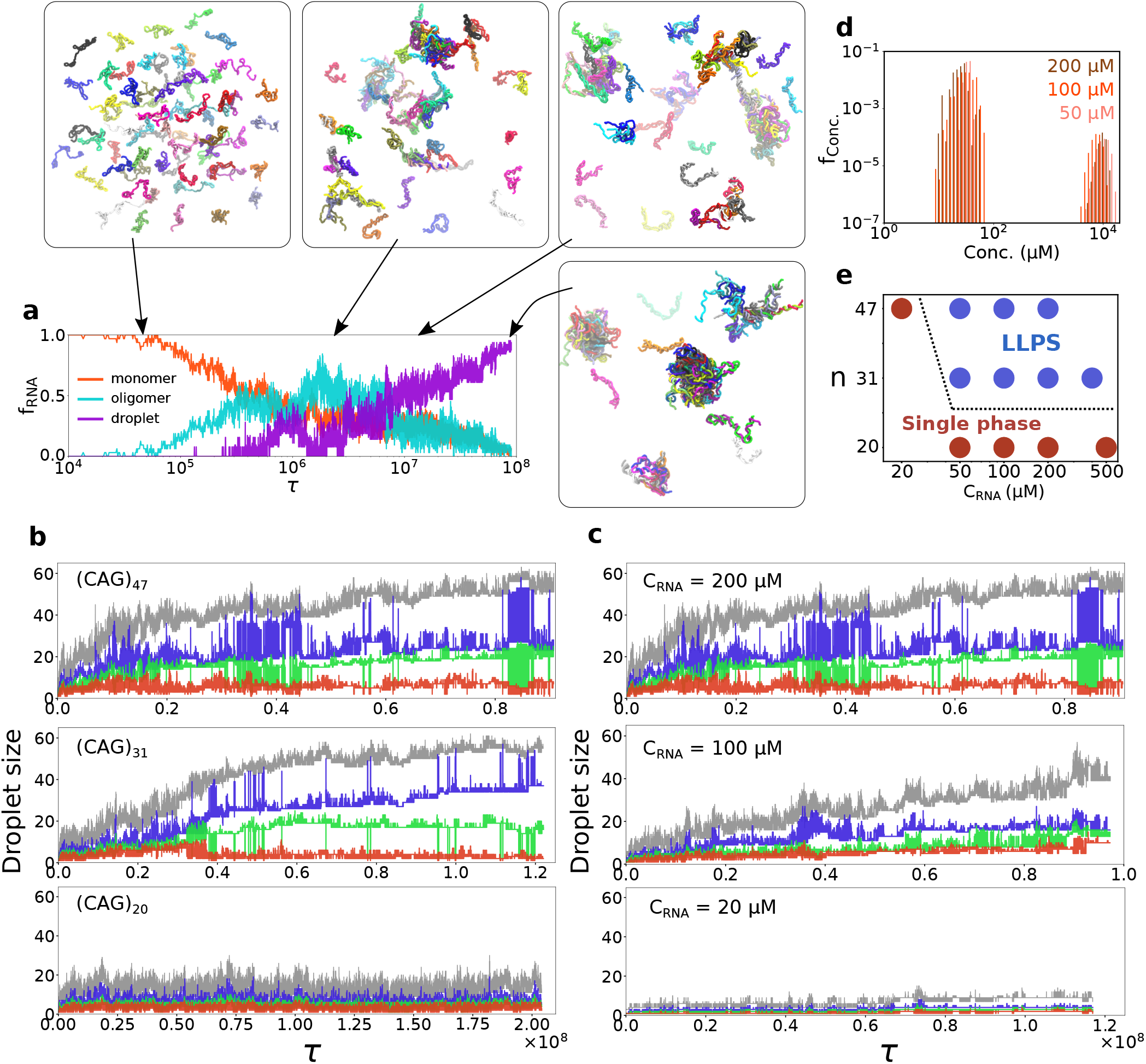
Liquid–liquid phase separation of CAG repeat RNAs. **a**, Fraction of (CAG)_47_ in monomers, oligomers (dimers, trimers and tetramers) and droplets as a function of time. (Oligomers have 2-4 chains, droplets have ≥ 5 chains, see more in Fig. 3**c**.) At *τ* = 0, the RNA chains exist as monomers, whose structural characteristics are described in the Supplementary Information (SI). Subsequently, oligomers form, which fuse together to form large droplets. Snapshots of (CAG)_47_ pictorially illustrate phase separation. **b**, Evolution of the three largest droplet sizes (blue, green and orange, respectively) for different *n*. The droplet size is defined as the number of RNA chains inside the droplet. The grey curve is the sum of the other three. (CAG)_47_ and (CAG)_31_ form several large droplets while (CAG)_20_ shows little sign of phase separation. **c**, Concentration dependence of (CAG)_47_ phase separation. From top to bottom, the RNA concentration decreases from 200 *µ*M to 20 *µ*M. At 200 *µ*M, very few monomers exist in the solution, as they all interact with each other to form stable droplets. At 20 *µ*M, stable condensate is not observed. **d**, Concentrations of the two phases showing the coexistence of low and high density phases of (CAG)_47_. The concentrations of the two (coexisting) phases are almost invariant, regardless of the initial concentrations, and are separated by almost 3 orders of magnitude. **e**, Putative phase diagram of (CAG)_n_. For *n* = 20, there is only one phase even at elevated RNA concentrations.

We then examined the effect of chain length *n* on the phase separation in (CAG)_n_. Fig. 1**b** shows that reduction of *n* from 47 to 30 has little effect on the phase transition behavior. However, if *n* is smaller than a critical number *n*^*^, the propensity to form liquid droplets ceases entirely. In the (CAG)_n_ system, *n*^*^ is somewhere between 20-30.

Next, we investigated the effect of RNA concentration, 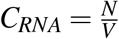, (*N* is the number of RNA molecules and *V* is the volume of the simulation box) on the condensate formation. We reduced (increased) *C*_*RNA*_ by increasing (decreasing) *V*, while keeping the number of chains fixed, *N* = 64. Fig. 1**c** shows that as *C*_*RNA*_ for (CAG)_47_ decreases, there is a decrease in the droplet size. At 200 *µ*M, almost all the molecules phase separate and enter the high density phase, leaving very few RNA chains as monomers in the solution. The two largest droplets (blue and green curves) each with ∼ 20-30 monomers frequently interact with each other. When *C*_*RNA*_ decreases to 50 and 100 *µ*M, only small droplets (< 20 monomers) form. The fluctuations in the size of these droplets are also much less pronounced, indicating that they are less likely to interact with each other and exchange monomers/oligomers.

We calculated the RNA concentrations of the coexisting low and high density phases using a procedure outlined in the Supporting Information (see also Fig. S2). The RNA concentration inside the droplets is enhanced by ∼50-200 fold compared to the initial concentrations (shown in Fig. 1**d**), in agreement with the experimental value (163-fold).^33^ Regardless of the initial *C*_*RNA*_, the concentrations of the two phases are relatively unchanged. Once the initial concentration of (CAG)_47_ reaches 20 *µ*M, which is lower than the concentration of the aqueous (or dispersed) phase, stable droplets do not form. At low concentrations, the RNA molecules mostly exist either as monomers or oligomers, and the system as a whole is a single liquid phase. Thus, for a fixed *n* that is larger than *n*^*^, there is a threshold value of *C*_*RNA*_, which has to be exceeded in order to form stable condensates. From the simulations at different *n* and *C*_*RNA*_, we determined the putative phase diagram for (CAG)_n_ (Fig. 1**e**). It shows that for *n*≤ 20, RNA molecules exist in a single phase. For *n >* 30, we predict coexistence of the high and low density phases, resulting from LLPS of the CAG repeat system.

### RNA structure changes dramatically in the condensates

We then characterized the RNA conformations inside the condensates containing multiple chains utilizing numerous quantities widely used in polymer physics. The conformational changes of (CAG)_n_ relative to the monomers is dramatic. (i) First, we calculate the mean distance between two nucleotides separated by *s* = | *i*− *j*| (*i* and *j* are the index of these nucleotides along the chain). The *R* (*s*) curve (shown in Fig. 2**a**) increases at large *s*, implying that the two ends are not in proximity, as in the isolated chain (see Fig. S3). (ii) Interestingly, *R* (*s*) versus *s* is independent of the droplet sizes. (Oligomers: 2-4 monomers, medium droplets: 5-10 monomers, large droplets: > 10 RNA molecules.) (iii) The bond orientational correlation, 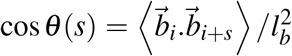 (*l*_*b*_ is the bond length), of the individual chains lacks periodicity inside the droplets, which is prominent in the monomers and oligomers (Fig. 2**b**). (iv) The end-to-end distance (*R*_*ee*_) distribution for chains inside the droplets shifts to higher values (Fig. 2**c**), signaling a disruption of the hairpin structures adopted by the isolated chains. Snapshots from the simulations (Fig. 2**e**) show that the RNA polymers populate extended conformations.

**Figure 2.**
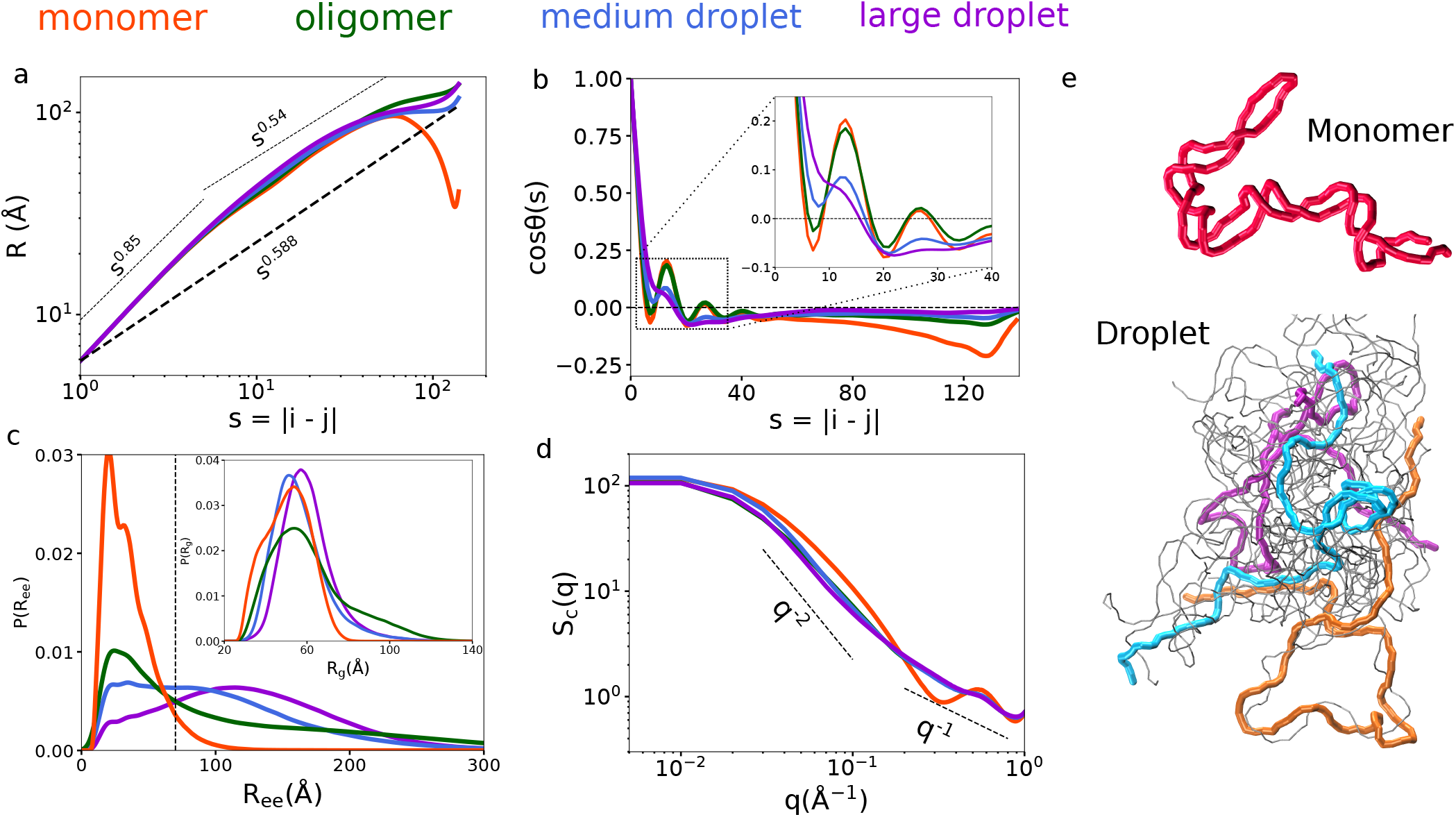
Comparison of the structures of (CAG)_47_ inside the condensates and in isolation. The colors indicate if the molecule is a monomer, or in different sized droplets. **a**, *R*(*s*) vs. *s* = | *i*− *j* |. The dashed line shows *R*(*s*) ∝ *s*^0.588^ for a self-avoiding polymer. **b**, Bond orientational correlation function, cos *θ* (*s*), as a function of *s*. Note the absence of periodicity (see inset for an enlarged view) in the large droplet (purple), which is prominent in the monomers and oligomers. **c**, Distributions of the end-to-end distance, *R*_*ee*_. The vertical dashed line is the average value for an ideal chain. The *R*_*ee*_ values inside the condensates are significantly larger compared to monomers, and the distribution could be fit using a broad Gaussian (see the inset in Fig. 4**a** for the fit). Distributions of the radius of gyration (*R*_*g*_) are in the inset. **d**, Form factors of RNA chains show that the RNA molecules inside the droplets are similar to an ideal polymer. **e**, Snapshots highlight the conformational differences between isolated chains (top) and chains inside the condensates (bottom).

The form factors (Eq. 5 in the SI), which characterize the overall shape of the polymer in the reciprocal space, show (Fig. 2**d**) that the chains inside the droplets are markedly different from the monomeric RNAs. For the chains in the condensates, at small scattering vector 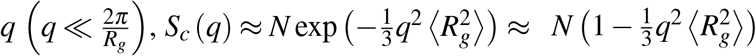 in the Guinier regime. At higher *q*, there is a crossover to a power law, *S* (*q*) ∼ *q*^−1*/ν*^, with *ν*≈ 0.5, suggesting that the RNAs may be characterized as ideal chains just as in polymer melts. This is because from the perspective of a single chain inside dense condensates, it is irrelevant whether the interactions arise intramolecularly or from other chains in the droplet. A similar reasoning led to the “Flory ideality hypothesis” proposed for polymer melts (see Fig. 3 for the analyses of intra- and intermolecular interactions). At small spatial length scale, or high *q, S* (*q*) ∼ *q*^−1^ is recovered.

**Figure 3.**
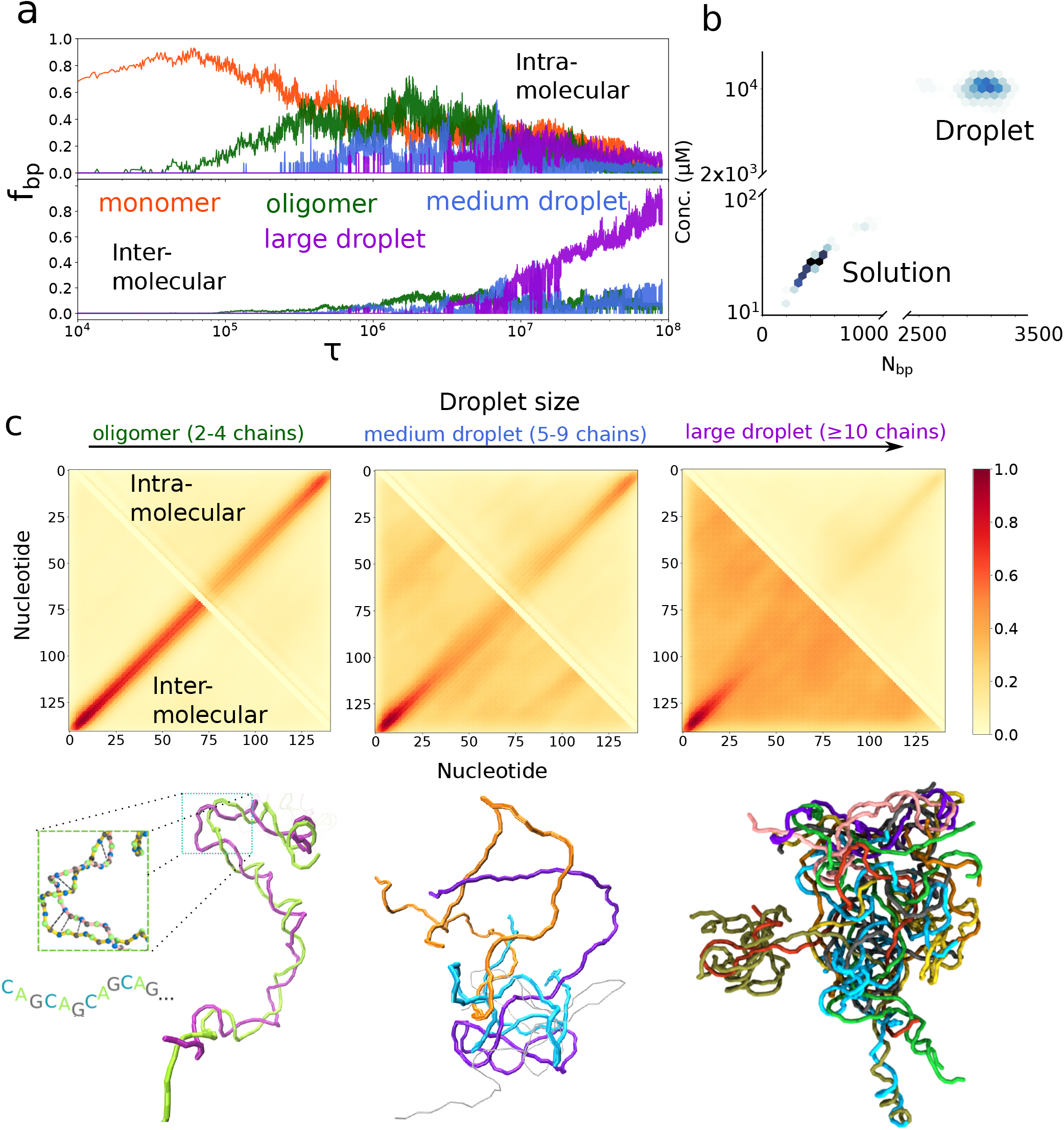
Intermolecular hydrogen bonding drives LLPS in RNA. **a**, Fraction of base pairs, *f*_*bp*_, between G and C nucleotides in (CAG)_47_ at 200 *µ*M (results for others are in Fig. S13). Decomposition into intra- and intermolecular contributions (top and bottom panels, respectively). **b**, Concentration vs. the number of base pair formed in the two coexisting phases. **c**, Probability of contact formation due to intra- (top) and intermolecular (bottom) interactions (color scale shown on the right). The contact between the two nucleotides is formed whenever the distance between them is less than 20 Å. Left to right correspond to oligomers, medium and large droplets, respectively. In oligomers, intermolecular interactions between the RNAs generate structures similar to RNA duplexes, with signals along the anti-diagonal (from the bottom left to the top right) in the contact map. In larger droplets, the specific interactions decrease, and non-specific interactions predominate. Representative snapshots for different droplets with different sizes are at the bottom.

### Intermolecular base-pairing drives RNA condensate formation

We calculated the fraction of formed base pairs (bps), 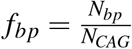 where *N*_*bp*_ is the number of bps and *N*_*CAG*_ is the number of CAG units in the simulations, to decipher the molecular details of the transition from low order oligomers to condensates (see SI for details). The results were decomposed into intra- and intermolecular bps (shown in Fig. 3**a**). Initially, the chains mostly adopt hairpin-like structures by forming exclusively intramolecular bps. Once small and medium sized droplets form, intermolecular interactions increase at the expense of intramolecular interactions. In order for RNA chains to form interchain bps, the C and G nucleotides from two chains have to satisfy both the distance and orientation criteria, thus requiring population of extended conformations (Fig. 2). In this process, the self-interactions are replaced by the intermolecular interactions without having to compensate for bending on short length scale, *s*.

We find that intermolecular base pair interaction is the driving force in the coalescence of the RNA repeats.^18, 33, 44, 45^ For short chains, (*n* = 20) or at low *C*_*RNA*_, the number of intermolecular interactions is small (Fig. S13). Thus, the majority of the molecules remain as monomers or form relatively small droplets with less than 8 chains. At high concentrations or for longer chains (*n* = 47 and *n* = 31), increase in the number of intermolecular bps leads to phase separation, resulting in the formation of two discrete phases with vastly different RNA densities. The number of bps in the high density phase is much higher than the low density phase (shown in Fig. 3**b**).

Intra- and intermolecular interactions depend on the droplet size (Fig. 3**c**). Unlike the self-interactions in the isolated RNA chain, which feature extensive WC base pairing along the anti-diagonal to form hairpin structures (Fig. S3), those inside the droplets are structurally more diverse. The propensity to form interactions along the chains increases, beyond those that form along the anti-diagonal. The total number of self-interactions of RNAs inside the droplets is smaller than in the monomers (Fig. 3**a**) due to the formation of extended conformations. Because the RNA conformations are significantly stretched inside the condensates, the interactions between different chains ought to increase. Intermolecular interactions between RNA molecules in droplets are also position-independent, reflecting the repeat nature of the sequence. However, for oligomers, specific 5’ to 3’ interactions still dominate, similar to what is found in a hybridized duplex.

### RNA conformations in the droplets are highly heterogeneous

Due to extensive intermolecular base pair formation inside the droplets, we expect the RNA mobility to be significantly hindered. Each RNA could be kinetically trapped, and only sample conformations around a local minimum in the free energy landscape, much like in a glass or a jammed system.^46^ Fig. 4 reveals that this is indeed the case. The distributions of *R*_*ee*_ (end-to-end distance) and *R*_*g*_ (radius of gyration) of the individual chains have large dispersion exceeding their usual values, although the histogram averaged over all the chains show behavior resembling that expected for ideal chains (red curves). Therefore, averaging over the ensemble of structures and over the number of RNA monomers in the droplet conceals the conformational heterogeneity. Some of the chains sample conformations with *R*_*ee*_ exceeding 200 Å, which is much greater than the dimensions expected for a random coil. As a consequence, there is a great degree of heterogeneity in the conformations that are sampled in the droplets. The unusual conformations that are accessed could arise because of the droplet is arrested in a non-ergodic phase with long overall relaxation times.

**Figure 4.**
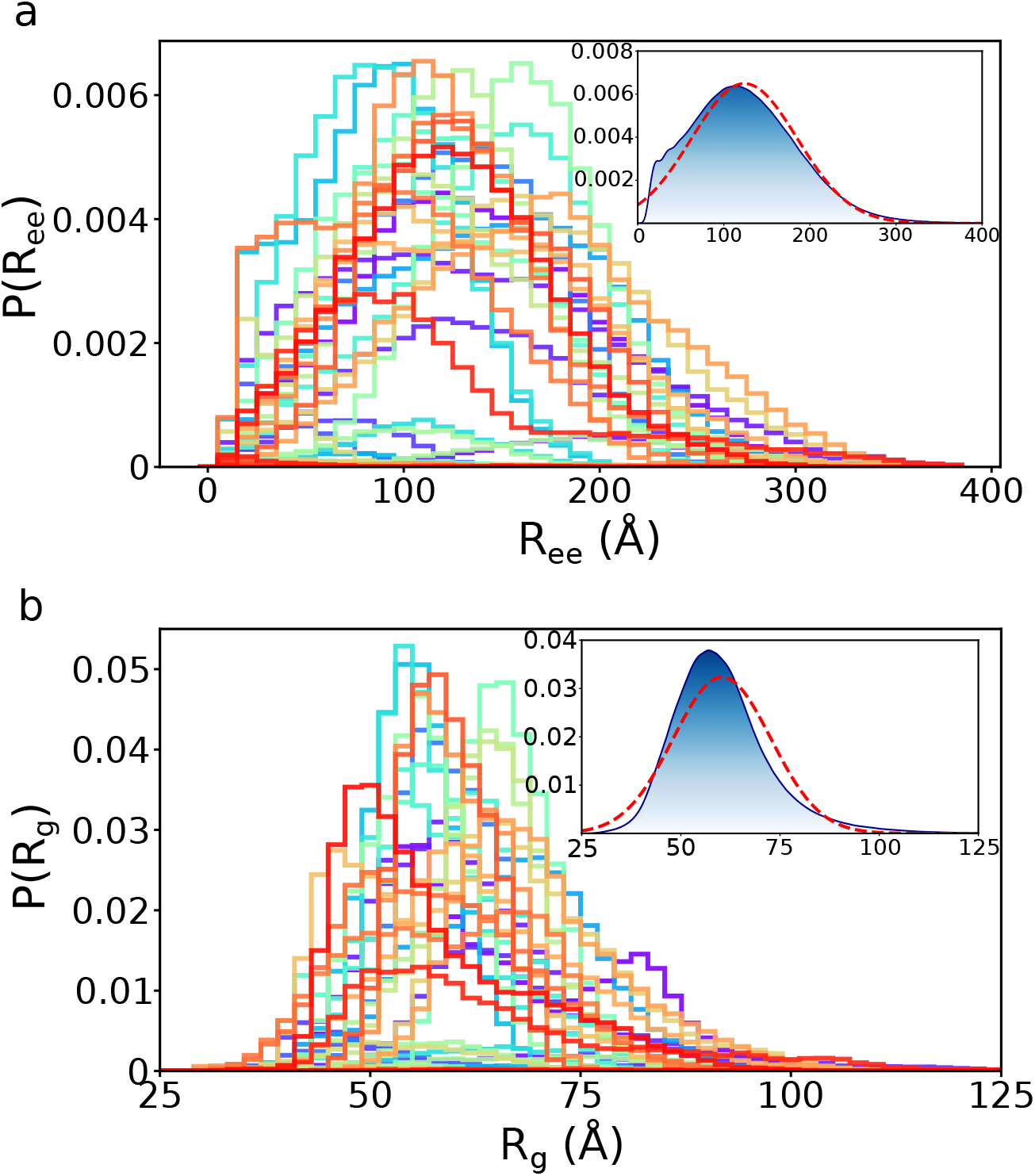
Structural heterogeneity of the RNA chains in large and dense (CAG)_47_ droplets. Distributions of *R*_*ee*_ **(a)** and *R*_*g*_ **(b)**, respectively, for each individual RNA chains. The insets show the average distributions, and the red dashed curves are the Gaussian fit to the data, indicating the behavior of chains inside large droplets is similar to ideal chains. Averaging over the monomers completely masks the remarkable heterogeneity of the monomer structures in the droplets.

### Condensates are dynamic

A hallmark of liquid–liquid phase separation is that the droplets (or foci) interact with one another, which could result in two (or more) droplets fusing to form a larger droplet by the Ostwald ripening mechanism. The reverse process in which a large droplet could disintegrate into smaller ones could also occur. Our simulations capture both these processes in the phase separation of (CAG)_n_. Fig. 5**b** shows the time evolution of each droplet size in the system, illustrating that the condensates formed in the simulations are dynamic, undergoing continuous growth, fusion and fission. The small droplets frequently interact and fuse with one another to form a larger droplet (snapshots shown in Fig. 5**a**, Movie S2). However, not all fusion events are successful. There are instances of failed events in which the two droplets come together, interact for a while but subsequently dissociate (Fig. 5**a**). Such fusion-fission processes occur frequently, suggesting that the droplets are dynamic.

**Figure 5.**
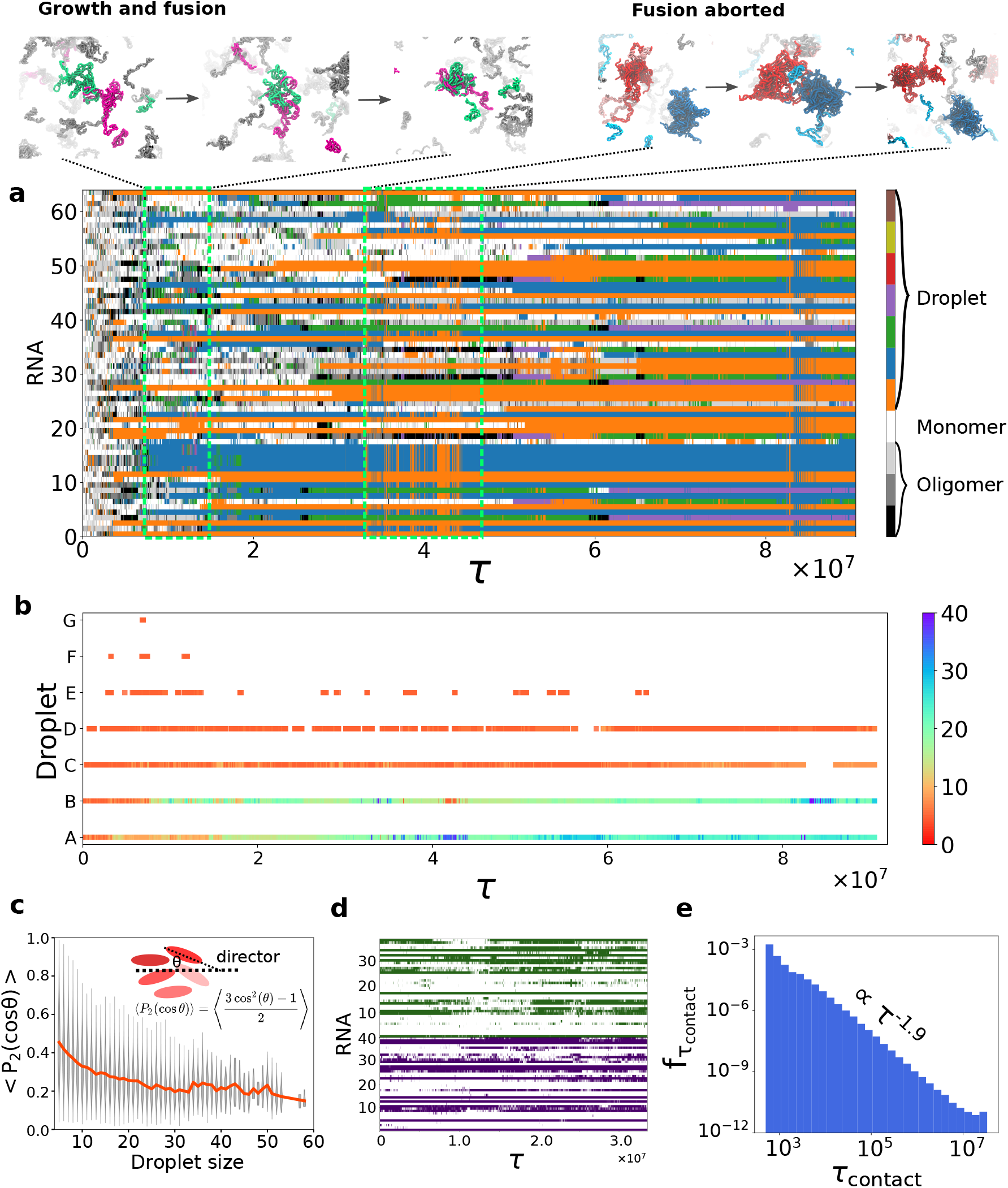
Condensate dynamics. **a**, Each line represents time evolution of a single chain. Colors encode the states of RNA. White is a monomer, oligomers are from grey to black. Seven droplets with different colors contain several RNA chains. Droplet growth and fusion shown on top left. Only a few chains are colored, the rest are shown as grey molecules. Aborted fusion event shown on the top right when the two droplets are in proximity and subsequently dissociate. See also Movie S2. **b**, Evolution of droplets with the color indicating the droplet size (scale on the right), with each row representing a unique droplet, corresponding to one color in **a. c**, RNAs in small droplets have high nematic order, < *P*_2_ cos(*θ*) >, which decreases as the droplet size increases (the orange curve shows average values). The grey lines show that there are substantial order parameter fluctuations. **d**, Contacts between one particular RNA with other chains inside the same droplet as a function of time. Color code: white - no contact; color - contact. Shown are two plots for two different chains in the same droplet. **e**, Frequency of contact lifetime between chains inside large droplets. The majority of contacts is transient, and the distribution of *τ*_*contact*_ follows a power law decay.

The dynamic nature of the droplets is also highlighted by the lack of persistent internal order, measured by the nematic order parameter (Fig. 5**c**). For small droplets, the internal RNA molecules need to position themselves in a preferred orientation, which would result in a loss of entropy. An extreme example is a dimer, in which the two strands are parallel to maximize the interaction energy (illustrated in Fig. 3**c**). However, in large droplets the RNA could form bp interactions with other chains without sacrificing orientational entropy, which results in a decrease in the nematic order parameter.

Fig. 5**a** shows exquisitely the dynamic nature of the condensate formation. Each row corresponds to a single chain, with the color denoting the droplet to which it is recruited. Each color represents a unique droplet in Fig. 5**b**. At the early stage (*τ* < 2 × 10^7^), most RNA molecules, at one instance or another, interact with almost all the droplets. Their residence times within a single droplet is relatively short. The monomers exchange with the bulk and are subsequently recruited by another (or the same) droplet. In contrast, at later stages, the chains are mostly restricted inside large droplets (orange and blue droplets in Fig. 5**a**). Despite the restricted movement, the interactions between the RNA chains inside large droplets are nevertheless transient, as illustrated in Fig. 5**d**, which shows the contact lifetime of one molecule with all other chains inside a single droplet. Although some of the contacts are stable most of them are intermittent and are disrupted over time. The lifetime of contacts decay following a power law (Fig. 5**e**). This is because the chains shift their internal positions frequently, and therefore interact with different chains at various times (see Movie S2). Due to the absence of droplet fusion and lack of monomer exchange as time progresses, we surmise that the large droplets in our simulations could be near the onset of coarsening into a rigid gel-like state, which is in accord with *in vitro* experiments.^33^

A major advantage of our simulations is that it is possible to describe in vivid details the behavior of the chains during the phase transition process where the monomers become part of the droplets. We found that there is no average or typical way a chain is integrated into a droplet, which underscores the importance of conformational and dynamic heterogeneity. Fig. 6 illustrates one possibility (out of a large number) of how a droplet could grow, and what happens to the RNA chains during the process. Fig. 6**a** shows the number of base pairs for a particular chain in the (CAG)_47_ simulation. As the chain enters the droplet, the number of base pairs it forms with other chains increases, while the number of intramolecular base pairs diminishes. The recruitment of the monomer to a droplet started from a dangling end or an overhang of the hairpin (panels **d** and **e**). The end of the chain then hybridized with other chains on the surface of the droplet, and gradually penetrated into the droplet interior. Interestingly, the penetration of the monomer into the droplet was associated with slippage events of the stem region of the monomer hairpin. It is possible that the nature of the repeat sequences makes this kind of integration energetically feasible, as the monomer could retain intra-base pairs that stabilize the stem on one side, while the other end hybridizes with other chains in the droplet. We wish to emphasize that the mechanism of incorporation is different for other chains. In addition, the droplet could grow by recruiting not only monomers but also oligomers and even smaller droplets (Fig. S9 and Movies S1, S2). The growth mechanism is, therefore, likely highly heterogeneous.

**Figure 6.**
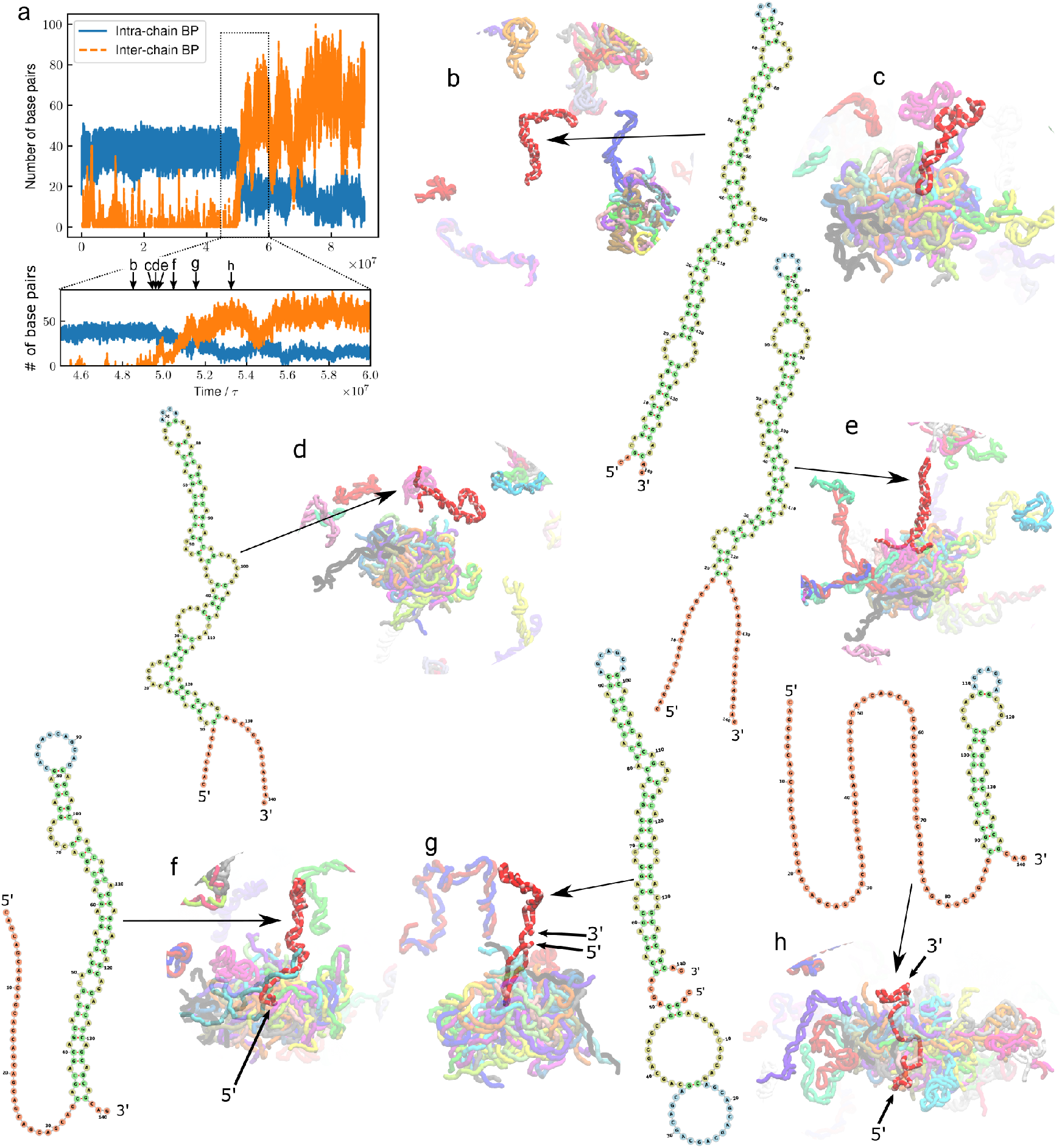
Visualization of the dynamics of monomer recruited into a droplet for (CAG)_47_. This is merely one scenario that illustrates the changes in the monomer dynamics as it is recruited into the droplet. See Movies S1 and S2 for additional scenarios. **a**, Number of base pairs of one specific chain as the recruitment process occurs. The base pairs are classified as either intra- (blue) or inter-chain (orange). In the bottom panel, the time is enlarged with arrows indicating time points of snapshots (**b**-**h**). (**b**-**h**), Snapshots taken from the monomer recruiting event, accompanied by the RNA secondary structure of the recruited chain. **b**, Initially, the chain (in red) is in the hairpin conformation. **c**, The chain approaches a droplet from the tip of the loop. The attempt to coalesce is unsuccessful, and the chain eventually dissociates from the droplet. **d**, The 5’ and 3’ ends open, creating overhangs. **e**, The opened ends start interacting with other chains on the surface of the droplet. **f**, The 5’ end starts expanding and hybridizes with another chain (cyan) in the droplet. The base pairs of the hairpin stem slips due to the nature of the repeat sequence. As a result, the 3’ end shortens. **g**, At this stage, the 5’ end has been entirely integrated into the droplet. The 3’ end appears at the periphery, and transiently forms intramolecular base pairs. **h**, The RNA chain completely penetrates into the droplet from one side to the other. The 3’ end still has a short stem that is stabilized by intra-chain base pairs. This residual structure remains for the rest of the simulation. The secondary-structure diagrams were generated with Forna^47^.

### Monomer dynamics inside the condensates is sluggish: Evidence of reptation

We quantified the dynamics of RNA movement using the mean squared displacement (MSD) of the RNA center-of-mass (Fig. 7**a**). The average MSDs for all the RNA chains increase linearly with time regardless of *C*_*RNA*_, implying that the overall motion is diffusive. However, the dynamics is highly heterogeneous upon undergoing LLPS. The RNA chains in the aqueous phase undergo normal diffusion, but the movement of the polymers in the droplets are significantly affected, especially in large droplets. The MSD exponents are less than unity, and decrease as the droplet size increases (Fig. 7**b**), which is suggestive of sub-diffusive behavior. Indeed, the slowing down of the RNA dynamics inside large droplets could signal the onset of formation of a gel-like state. Given sufficient time, the large droplets could potentially undergo subsequent maturation through a sol-gel transition.^20, 46, 48–56^

**Figure 7.**
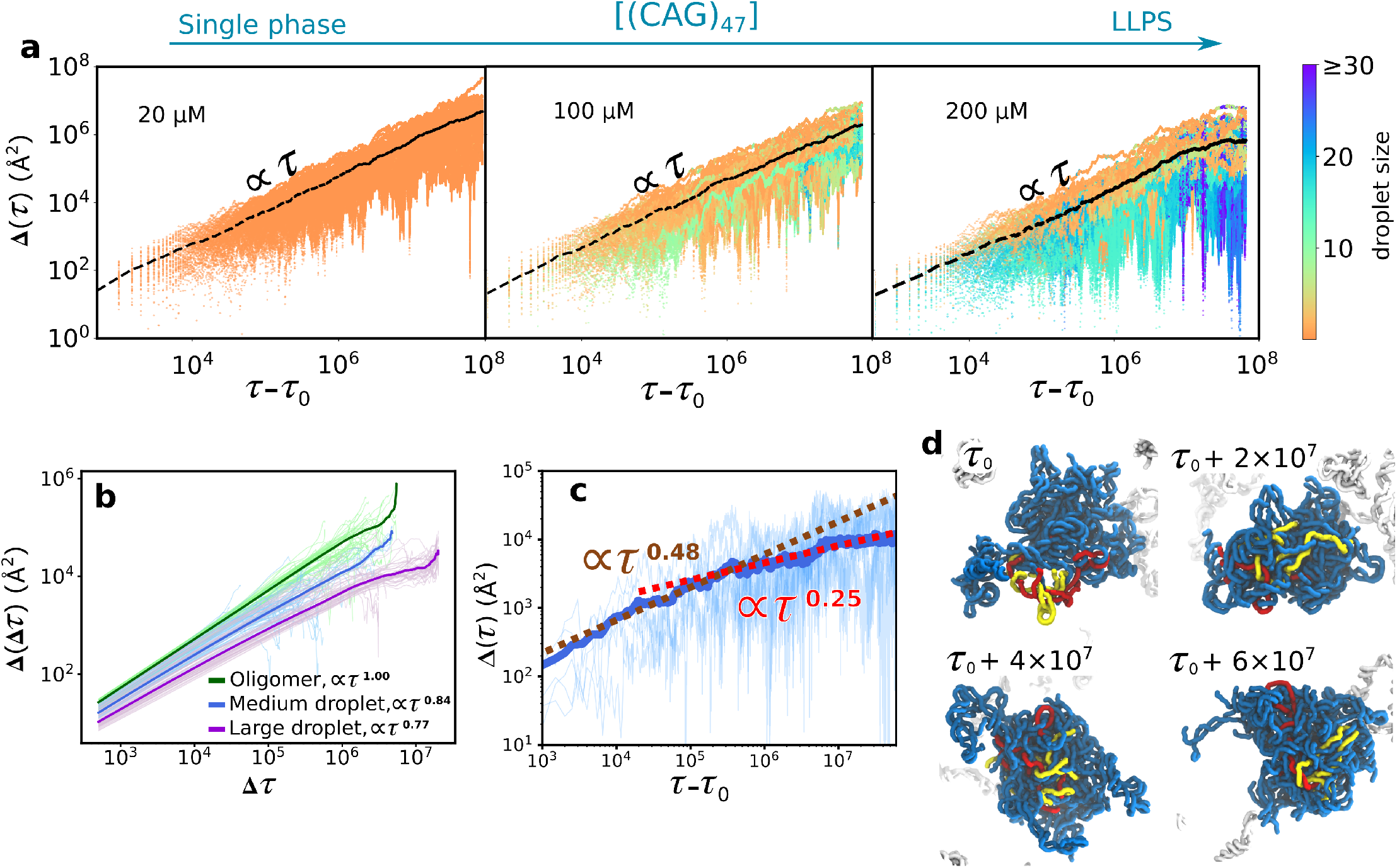
Dynamics of RNA chains in (CAG)_47_ condensates is heterogeneous. **a**, Mean squared displacement (MSD) of the RNA center-of-mass at different concentrations, increasing from left to right, shown in a log scale. Results are shown for individual RNA molecules with color denoting the droplet size. Averaged values are plotted as black lines. The majority of fast moving RNA chains is in oligomeric and monomeric states, while sluggish movement arises from RNA chains inside large droplets. **b**, Time-averaged MSD calculated for RNA chains inside oligomers, medium, and large droplets. The RNA movement in oligomers is diffusive, while for medium and large droplets, they undergo subdiffusive motion. Results for individual RNA molecules are plotted in pale color lines, and the average is shown as solid lines. **c**, Nucleotide MSD in a large viscous droplet. Solid line is the averaged value. In the intermediate regime (*τ* − *τ*_*o*_ *<* 10^5^, where *τ*_*o*_ is the initial time when the droplet forms), the MSD exponent is ≈0.5, indicating that the chains are jammed. There is a change in slope at *τ* − *τ*_*o*_ ≈10^5^, where Δ ∝ *τ*^1*/*4^. This suggests that the movement of connected nucleotides is reminiscent of reptation, which implies that chain dynamics is mainly restricted along the contour length of the RNA due to entanglement. The reptation dynamics is most clearly illustrated in Movie S3. **d**, Representative snapshots show movement of individual chains inside large droplets.

To probe the jamming dynamics of RNA chains inside large droplets, we calculated the MSDs for each nucleotide in the RNA chain (Fig. 7**c**). The average values (the solid line in Fig. 7**c**) show that the MSD scales as Δ(*τ*) ∝ *τ*^1*/*4^ at long times. In monomers, we find that Δ(*τ*) ∝ *τ*^0.78^ (Fig. S10). The *τ*^1*/*4^ behavior, which was first predicted theoretically^43^ and observed in simulations of polymer melts^43, 57, 58^, is suggestive of reptation-like movement. The motion of these nucleotides is physically constrained along the contour length of the RNA, reminiscent of the reptation mechanism in which the chain movement is thought to occur in a low (one) dimensional “tube” generated by topological constraints imposed by other chains (see Movie S3 for illustration). In order to exhibit reptation-like dynamics, the RNAs should form an intricate and entangled network of interactions, which arises due to intermolecular base pair interactions. Interestingly, the formation of such a network has recently been demonstrated in AU-rich mRNAs.^27, 59^ The network of intermolecular interactions between the RNAs, therefore, plays a crucial role in dictating the slow heterogeneous dynamics of RNAs inside the droplets.

## Discussion and Conclusion

We developed a minimal coarse-grained SIS model that captures many of the observed features in the LLPS of repeat RNA sequences. In accord with experiments, we find that long (CAG)_n_ repeats (*n* = 47, 31) phase separate at relatively low concentrations, whereas short chains (*n* = 20) remain in a single liquid phase even at elevated RNA concentrations. It is worth noting that there is only a single parameter in the SIS model that was determined using the structures of (CAG)_2_ duplex. In probing condensate formation in CAG repeats, the SIS model is essentially parameter-free, and hence is transferable to probe other RNA sequences as well. As an example, we show in Fig. S8 that the tendency to undergo LLPS in one scrambled sequence is diminished compared to the CAG repeat sequence, which is in qualitative agreement with experiments.^33^

### Recruitment of monomers to condensates involves unwinding of hairpin-like structure

Due to the high GC content of the sequence, the monomeric RNAs adopt an ensemble of hairpin-like structures with a small end-to-end distance (Fig. S3), as is the case in a number of RNA molecules.^42, 60, 61^ In sharp contrast, RNA chains inside the condensates are extended with large 5’-3’ distances. In order to form intermolecular interactions with other chains, the RNA monomer has to unwind, exposing the base to form WC base pairs (or *π* − *π* stacking and non-canonical base pair depending on the sequence^24, 27, 34^). We expect this finding to be a general feature of not only repeat RNAs, but also in any binary biomolecular system that undergoes phase separation (with solvent being another component), as was shown previously in oligomer formation in a fragment of A*β* peptides.^62^

### Topological constraints, conformational and dynamical heterogeneity

As a result of a large number of intermolecular interactions between the RNA chains within the condensates, a given RNA molecule is topologically constrained by the neighboring chains. The local environment of individual RNA chains differs greatly, which results in wide variability in the individual distributions of various shape parameters (Fig. S15), even though the average distribution could be explained using polymer theory. Because of such topological constraints, the dynamics of RNA chains is also strongly affected. The mobility of the RNA chains is greatly diminished even though diffusion of the system as a whole is liquid-like.^63, 64^ In large droplets in which long RNA polymers are tightly packed with elevated density and viscosity, the movement of RNA molecules is reminiscent of reptation in polymer melts. RNA chains slither along their contour length due to translational inhibition in other dimensions. Strong entanglement in large droplets could further facilitate the maturation process, eventually driving the system to a gel-like state at long times under *in vitro* conditions. Under cellular conditions, it is likely that active processes (ATP binding and/or hydrolysis) regulate the formation of gel-like states by remodeling the RNA structures.^65–68^ For instance, the recruitment of additional protein clients^29^ or the presence of ATP-consuming enzymes (for example, helicases or RNA chaperones) in the nucleoplasm that reorganize the RNA base pairings^33, 68^ could play a role in maintaining the condensates in a liquid-like state.

### Limitations of the study

The simulations, using the SIS model without any parameter to fit the experiments, explains the findings on condensate formation in the repeat RNA sequence. However, the SIS model has limitations.

1. Counter ions, which play an important role in driving condensate formation,^33, 69^ are not included explicitly. We assume that the cumulative effects of the Tris buffer, the added monovalent and divalent ions used in the experiments effectively screen the electrostatic repulsions between the nucleotides. It has been shown by Oosawa–Manning and confirmed by experiments that the polyelectrolyte charge is neutralized by nearly 88% whenever divalent ions are present (compared to 76% for monovalent ions alone).^70–72^ Under these conditions, the electrostatic repulsion between the phosphate groups is effectively 1% of its original magnitude, justifying our assumption of complete neutralization of RNA charge by counterions. Our simulations are valid at high ion concentrations at which the charge on the phosphate is negligible. The impact of ions, especially divalent cations, has to be examined, possibly using the theory proposed recently,^73^ for a more complete theoretical understanding of the RNA phase separation.
2. Our simulations only consider the canonical Watson–Crick (WC) bp formation, and completely neglect the possibility that non-canonical base pairs can form. This assumption is rooted in the observation that only 20-30% of nucleotides form non-canonical bps in the RNA structures.^74, 75^ Another reason is that the difference between these types of bps mostly arises from the alternative conformations of the base, sugar and phosphate groups within the same nucleotide. Since the SIS model represents a nucleotide by a single bead, any distinction at the single nucleotide resolution cannot be modeled explicitly. In order to take into account the possibility of non-canonical base pair formation, one needs to utilize the more detailed three-interaction-site model in which a nucleotide is represented by three beads for the base, sugar and phosphate groups. Incorporation of such non-canonical bps in the simulations could introduce alternative conformations and intermediate structures during the phase transition process (Fig. S5). However, we anticipate that such modifications would not significantly change the outcomes of the simulations.^76^
3. It is unclear whether the *in silico*-generated condensates adopt gel-like states that has been suggested using the FRAP recovery experiments, which we should point out are not always accurate. We did provide evidence that the condensates start the transition from liquid-like to gel-like at long times. For instance, as time progresses, we have shown that the droplets fail to fuse once they are close and the monomers inside droplets stop exchanging with the diluted phase. These criteria are qualitatively similar to how gel-like or liquid-like nature of the condensates are characterized in experiments. Our simulations, therefore, suggest that the condensates first form via LLPS, and then subsequently coarsen into “gels”, which could be the state observed in the Jain–Vale experiment. The timescale of the maturation process is likely to be long compared to the simulation timescale presented here. Because of the difficulty in distinguishing between the sol state with very slow dynamics and a gel-like state, material properties (by applying shear for example) have to be determined before further characterization of the RNA condensates.^46^

### Final remarks

We have used molecular simulations to probe condensate formation in repeat RNA sequences, and have uncovered general concepts that are difficult to obtain using experiments alone. Our study establishes that in the process of self-assembly of low complexity RNA sequences into droplets, RNA chain makes a transition from hairpin-like structures to extended conformations in order to maximize the number of intermolecular base pair interactions. The droplets are stabilized by a network of soft intermolecular interactions, which repeatedly break and reform as the chains move within the confines of the droplets. The dense network, which is most prominent in large high density droplets, restricts the mobility of the chain mostly along the contour. As a result, motion occurs by a process that is similar to snake-like movement envisioned in the context of polymer melts. Taken together, our results show that the structural heterogeneity and the pinning of RNA chains by their neighbors makes the dynamics sufficiently sluggish that the droplet may be in a non-ergodic phase.

The entanglement of RNA chains arises in part because the CXG sequence could engage in both intra- and intermolecular WC base pair formation. Because such specific interactions are likely not possible in poly-rU sequences,^77^ which also form droplets stabilized possibly by *π* − *π* stacking interactions, we believe that the only requirement for droplet formation in low complexity RNA sequences is the presence of many weak inter-chain interactions. This would suggest that the length dependence for condensate formation is likely to be different in poly-rU sequences than in CAG or CUG polymers. Future studies are needed to provide definite answers to the effect of base stacking and non-canonical base pair formation on RNA phase separation. Finally, it is worth emphasizing that our computational framework is not only well suited to examine the mechanism of self-assembly in other RNA sequences but also is general enough to account for ion effects and other cellular factors.

## Data availability

All data are included in the article and/or the SI. The raw data is available from the corresponding author upon reasonable request.

## Code availability

The codes to perform simulations and analyses are available at GitHub (https://github.com/tienhungf91/RNA_llps).

## Acknowledgements

We are indebted to Anne D. Bowen at the Visualization Laboratory (Vislab), Texas Advanced Computing Center, for generating the videos. We are grateful to Hiranmay Maity, Sua Myong, Mauro Mugnai, Sumit Sinha and Ryota Takaki for stimulating discussions and critical reading of the manuscript. This work was supported by National Science Foundation Grant (CHE 19-00093) and the Welch Foundation Grant (F-0019) through the Collie–Welch chair. We are thankful to the Texas Advanced Computing Center for providing computational resources.

## Author contributions statement

H.T.N. and D.T. conceived and designed research, H.T.N. conducted research, H.T.N., N.H. and D.T. analysed the results and wrote the manuscript.

## Notes

### Competing Interest Statement

The authors have declared no competing interest.

