## Supplementary Information for "Condensates in RNA Repeat Sequences are Heterogeneously Organized and Exhibit Reptation-like Dynamics"

### Supplementary Methods

#### RNA energy function

In order to describe the organization and dynamics of RNA condensates, we develop a new coarse-grained model that is sufficiently simple to simulate multiple chains while retaining sequence information. Such an approach is needed to investigate the mechanism of RNA condensation in arbitrary sequences. Because condensate formation requires multiple simulations involving many chains for sufficiently long times, we resorted to a low resolution coarse-grained description for RNA. We have shown previously that simulations based on related models are efficacious in predicting the outcomes of single molecule pulling experiments.[1,2]

We introduce the Single Interaction Site (SIS) model in which each nucleotide is represented by a single bead (Fig. S1c). The energy function for an isolated RNA chain has the following form:

$$U = U_{BA} + U_{EV} + U_{HB}, \tag{1}$$

where $U_{BA}$ accounts for chain connectivity, consisting of bond and angle restraints between the connected beads. We used harmonic potentials to keep the bond lengths and the angles close to the A-form helix: $U_{BA} = \frac{1}{2}\sum_i k_{bond}\left(r_i - r_o\right)^2 + \frac{1}{2}\sum_j k_{angle}\left(\alpha_j - \alpha_o\right)^2$ where $k_{bond} = 15.0$ kcal/mol.Å$^2$, $k_{angle} = 10.0$ kcal/mol.radian$^2$, $r_o = 5.9$ Å, $\alpha_o = 2.618$ rad. In the simulations, the angles fluctuate due to soft restraints, and deviate from the values in the A-form helix.

The excluded volume interaction $U_{EV}$ is given by the Weeks–Chandler–Andersen potential:

$$U_{EV} = \sum_{i,j} \Theta\left(\sigma - r_{ij}\right)\varepsilon\left[\left(\frac{\sigma}{r_{ij}}\right)^{12} - 2\left(\frac{\sigma}{r_{ij}}\right)^{6} + 1\right], \tag{2}$$

where $\Theta$ is the Heavyside step function, $\sigma = 10.0$ Å, and $\varepsilon = 2.0$ kcal/mol. The excluded term is only computed between two beads that are not involved in the bond or angle restraints, and are separated at least by two other beads (nucleotides) along the chain.

Hydrogen bond term $U_{HB}$ is a many-body and short-ranged potential that mimics the canonical Watson–Crick base-pairing between A-U and G-C. The hydrogen bonding potential between the two beads $i, j$ is taken to be,

$$U_{HB} = U^o_{bp} \exp\left[U_{hb,bond} + U_{hb,angle} + U_{hb,dihedral}\right], \tag{3a}$$

$$U_{hb,bond} = -k_r\left(r_{ij} - r_{hb,o}\right)^2, \tag{3b}$$

$$U_{hb,angle} = -k_\theta\left(\theta_{i,j,j-1} - \theta_1\right)^2 - k_\theta\left(\theta_{i-1,i,j} - \theta_1\right)^2 - k_\theta\left(\theta_{i,j,j+1} - \theta_2\right)^2 - k_\theta\left(\theta_{i+1,i,j} - \theta_2\right)^2, \tag{3c}$$

$$U_{hb,dihedral} = -k_\phi\left[1 + \cos\left(\phi_{j-1,j,i,i-1} + \phi_1\right)\right] - k_\phi\left[1 + \cos\left(\phi_{j+1,j,i,i+1} + \phi_2\right)\right], \tag{3d}$$

where $\theta_{a,b,c}$ is the angle formed between beads $a$, $b$ and $c$; $\phi_{a,b,c,d}$ is the dihedral angle formed between beads $a$, $b$, $c$ and $d$; $r_{hb,o} = 13.8$ Å, $k_r = 3.0$ Å$^{-2}$, $k_\theta = 1.5$ rad$^{-2}$, $k_\phi = 0.5$, $\theta_1 = 1.8326$ rad, $\theta_2 = 0.9425$ rad, $\phi_1 = 1.8326$ rad, $\phi_2 = 1.1345$ rad.

The functional form of $U_{HB}$ is designed to capture both the Watson–Crick base-pairing, and the helical nature of the A-form RNA at the single-nucleotide resolution. The only unknown parameter, the strength of the hydrogen bond interactions, is adjusted to reproduce the structure of a small CAG repeat sequence[3] ($U^o_{bp} = -5.0$ kcal/mol for GC base pair, equivalently $\approx -1.67$ kcal/mol per hydrogen bond, Fig. S1).

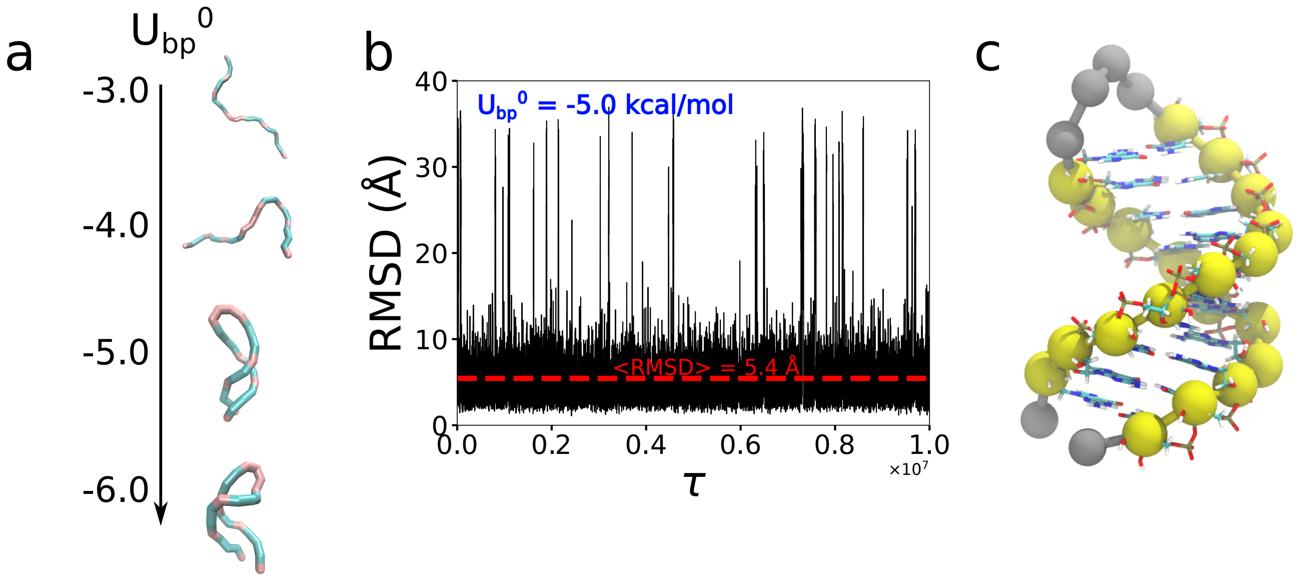


**Figure S1.** Calibration of the base pair interaction strength $U^o_{bp}$. **a**, Structural dependence of a small CAG repeat sequence (AGGCAGCAGCCAAAAGGCAGCAGCCA) on $U^o_{bp}$. The sequence we chose to calibrate $U^o_{bp}$ is almost identical to the X-ray structure (PDB 3NJ6),[3] except with the addition of an AAAA tetraloop and two terminal A nucleotides. The sequence adopts extended conformations for small $U^o_{bp}$ ($U^o_{bp} < 4.5$ kcal/mol), and folds into hairpin conformations in base pair interaction range, $4.5 < U^o_{bp} < 6.0$ kcal/mol. We set $U^o_{bp} = -5.0$ kcal/mol. **b**, Root mean squared deviation (RMSD) between the simulations and the X-ray crystal structure, shown for $U^o_{bp} = -5.0$ kcal/mol. The averaged value for RMSD is around 5 Å, which is reasonable given the coarse-grained nature of the model. **c**, Superposition of the simulated structure (yellow and grey beads) onto the X-ray structure for the lowest RMSD (around 1 Å).

**Inter-chain interactions:** Interactions between different chains also involve excluded volume and hydrogen bonding interactions, which are treated identically as in intramolecular interactions. Thus, the SIS model when used to simulate multiple RNA chains does not distinguish between intra- and inter-molecular base pair interactions. A nucleotide could form WC base-pair interactions with others either within the same chain or in a different chain. Therefore, the SIS model is a general enough for studying condensate formation in a generic RNA, provided that a single bead resolution suffices. As always, the validity of the model, like all others, can only be assessed by comparisons to experiments.

**Counterion effects:** Although that counter ions play an important role in driving RNA condensate formation (see panel h in the Extended Data Figure 1 in Ref.[4]), we do not include them explicitly in the current version of the model. One reason for neglecting them is that the charge on the phosphate ions has to be effectively neutralized by counterions in order for them to coalesce into droplets. Upon counterion condensation, electrostatic repulsion between phosphate groups is greatly diminished. A similar situation occurs in Ψ-condensation, where it has been shown that roughly 90% of the DNA charge is neutralized in order for the DNA condenses.[5,6] The electrostatic repulsion between phosphate groups is around 1% of its original magnitude. Thus, it is not unreasonable to assume that the effective charge on the phosphate group is sufficiently small that it be neglected. As a result, the simulations reported here are valid only at high monovalent (for example $Na^+$) and divalent ($Mg^{2+}$) concentrations. Future works, using a recently proposed theory,[7] could incorporate ion effects more precisely in order to calculate the ion-dependent phase diagram.

#### Simulation details and analyses

**Simulation details**. We placed 64 $(CAG)_n$ ($n$ = 20, 31 or 47) molecules evenly spaced in a cubic box, whose size varied from 50-200 nm depending on the RNA concentration. The initial conformations of the RNAs are significantly expanded to minimize possible conformational biases. The initial conditions used in the simulations are a mimic of the experimental protocol in which the RNA molecules are denatured at an elevated temperature for a certain duration, and then the temperature is decreased to room temperature. In order to ensure that our results are not affected by initial conditions, we also performed simulations where the temperature is decreased slowly starting from a high temperature. We observed little difference in the phase behavior (see details below), which shows that the results do not depend on the initial conditions as long as the RNA molecules are unfolded. Simulations were performed on GPU using the custom OpenMM code to speed up sampling of the conformational space.[8] We used low-friction Langevin dynamics in which the viscosity of water was reduced by a factor of 100 in order to further speed up the sampling efficiency.[9] Even using a the simple SIS model for RNA, the simulations are computationally intensive because there are many chains in the system. Simulations were conducted for ~ 100 days for each trajectory using NVIDIA Quadro RTX 5000 graphics card on the Frontera supercomputer (Texas Advanced Computing Center). Snapshots were recorded every 10,000 steps, which were subsequently used to calculate several quantities of interest.

**Clustering**. For each snapshot, we grouped the monomers that are within 20 Å from each other to the same cluster. A monomer belongs to a cluster if the distance between any of its nucleotide to any nucleotide in the cluster is less than 20 Å. For comparison, the equilibrium base pairing distance in the SIS model is $r_{hb,o} = 13.8$ Å. Thus, the cutoff distance for clustering is about $\approx 40\%$ larger than $r_{hb,o}$.

**Form factor.** The form factor of RNAs was calculated as:

$$S_c = \left\langle \left| \sum_k \exp\left(i\mathbf{q}.\mathbf{r_k}\right) \right|^2 \right\rangle_N , \tag{4}$$

where the brackets denote averaging over the chains at different orientation.

**Radius of gyration and shape parameters.** $R_g$ and shape parameters were calculated using the gyration tensor, with each element is defined as:

$$S_{xy} = \frac{1}{2N^2} \sum_i \sum_j \left(x_i - x_j\right)\left(y_i - y_j\right). \tag{5}$$

The radius of gyration, $R_g^2$, is then calculated by summing over the three eigenvalues of the gyration tensor $R_g^2 = \lambda_1^2 + \lambda_2^2 + \lambda_3^2$. To characterize the shape of RNAs, we calculated the relative shape anisotropy $\kappa^2$ and the shape parameter S using

$$\kappa^2 = \frac{3}{2} \frac{\lambda_1^4 + \lambda_2^4 + \lambda_3^4}{\left(\lambda_1^2 + \lambda_2^2 + \lambda_3^2\right)^2} - \frac{1}{2}, \tag{6}$$

$$S = \prod_{i=1,2,3} \frac{\lambda_1^2 - \bar{\lambda^2}}{\bar{\lambda^2}}, \tag{7}$$

where $\bar{\lambda^2} = \frac{1}{3}\left(\lambda_1^2 + \lambda_2^2 + \lambda_3^2\right)$. $\kappa^2$ is bounded between 0 and 1; $\kappa^2 = 0$ implies that the molecule is perfectly spherical, and $\kappa^2 = 1$ if every point lies on a straight line. $S$ satisfies $-1/4 \leq S \leq 2$. Negative values of $S$ correspond to oblate ellipsoids, while prolate ellipsoids have positive $S$.

**Concentrations of the two phases.** To determine the concentration inside a droplet, we first calculate the volume of a droplet. We assume the droplet is an ellipsoid with the three effective radii, characterized by the three eigenvalues of the gyration tensor. Thus, the volume of the i$^{\text{th}}$ droplet is,

$$V_i = \frac{4}{3}\pi abc = 4\pi\sqrt{3}\lambda_1\lambda_2\lambda_3. \tag{8}$$

Note that the gyration tensor is calculated for the whole droplet, and not individual RNA molecule as the previous section. The concentration inside the droplets is computed by averaging all the concentrations inside medium and large droplets:

$$C_{droplet} = \bar{C_i}, \tag{9}$$

with $C_i = \frac{N_i}{V_i}$ is the concentration of the i$^{\text{th}}$ droplet with $N_i$ RNA molecules and volume $V_i$. We calculate $C_i$ only for $N_i \geq 5$.

The concentration of the aqueous solution, consisting of monomers and oligomers outside the droplets, is then calculated using:

$$C_{solution} = \frac{N - \sum_i N_i}{V - \sum_i V_i}. \tag{10}$$

**Mean squared displacement.** The mean squared displacement (MSD) of the center-of-mass of a chain is given by,

$$\Delta(\tau - \tau_o) = \left\langle \left(\vec{r}_i(\tau) - \vec{r}_i(\tau_o)\right)^2 \right\rangle_N, \tag{11}$$

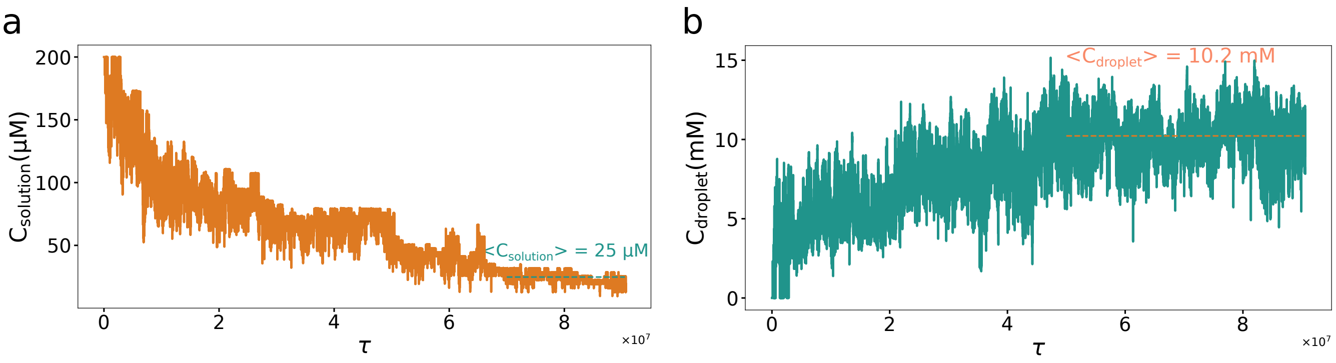


**Figure S2.** Determination of the concentrations of the two co-existing phases. Results are shown for $(CAG)_{47}$. Time dependent changes in the concentrations in the aqueous phase is on the left and for the droplet is on the right. The plateau values near the end are used to calculate the concentrations at which the two phases coexist.

where $\tau_o$ is the initial time.

The time-averaged MSD is calculated using:

$$\Delta(\Delta\tau) = \left\langle (\vec{r}_i(\tau+\Delta\tau) - \vec{r}_i(\tau))^2 \right\rangle_{\tau,N}. \tag{12}$$

We also computed the MSD of individual nucleotides to probe the dynamics of the RNA chains in large droplets (Fig. 7c in the main text). For this purpose, we chose the largest droplet formed in the simulations. Before computing the nucleotide MSD, the position of the droplet in each snapshot was sequentially superimposed to the one in the previous time frame, starting from the instance the droplet is formed ($\tau_o$). This procedure ensures that the MSD accounts solely for the dynamics of chains within the droplet, but not for collective translational and rotational motion of the entire droplet in the simulation box. The MSD was then calculated in the same way using Eq. 11 by averaging over all the nucleotides except for 10 nucleotides at both the $5'$ and $3'$ ends. Note that $\tau_o$ depends on when the droplet forms.

**Criterion for base pair formation.** To assess if a base pair (bp) is formed between G and C, we rely on the base-pair energy $U_{HB}$ defined in Eq. 3. If $U_{HB} < -3k_BT$, where $k_BT$ is the thermal energy, between two nucleotides, then we consider that they form a base pair. Since the energy function consists of several terms that depend on the distance, angles, and dihedral angles around the two nucleotides (Eq. 3b-d), the energy-based definition naturally captures the geometric criteria that base pairs satisfy in the ideal A-form RNA.

#### Supplementary Results and Discussions

##### Isolated repeat RNAs form hairpin-like structures

We first characterized the structural ensembles of an isolated $(CAG)_n$, which serves as a reference when comparisons to RNAs within the condensates are made. Interestingly, the mean end-to-end distance, $R_{ee}$, is independent of $n$ (Fig. S3**a**), which accords well with experiments and theoretical predictions for mRNA and long non-coding RNA.[10–13] The peak values in the distributions of $R_{ee}$ is around 35 Å. The $R_{ee}$ distributions for the three RNA chains with different lengths deviate from the Gaussian distribution for ideal chains, exhibiting long tails. This is somewhat surprising because the radius of gyration $R_g$ distribution (shown in Fig. S3**b**) shows little deviation from the ideal chain behavior.

Unstructured RNAs (poly-rA or poly-rC) that do not favor base pair formation follow the expected $R_{ee}$ distribution for a random coil.[13,14] In contrast, CAG repeats form intramolecular base pair interactions, bringing the 5' and 3' ends to proximity. Experiments have reported that $(CAG)_n$ repeat sequences with small $n$ could form stable hairpins containing GC base pairs (bps) with A:A mismatches.[3,15] Formation of GC bps in longer repeats also requires the chain to fold upon itself. Folding would be favored if the overall free energy gain by forming GC bps compensates for the chain bending penalty. For the CAG repeat sequences, extensive GC bp formation could mitigate the unfavorable interactions, leading to the formation of hairpin-like structures.[3,16] The probabilistic contact map (Fig. S3**d**) shows that the majority of interactions are along the anti-diagonal, consistent with the formation of a hairpin structure. The two ends have the highest probability of being in proximity. Representative snapshots from the simulations, shown in Fig. S3**a**, confirm that the RNA monomer mostly samples a set of hairpin-like structures.

It is worth emphasizing that we did not adjust any parameter in the SIS model to constrain $(CAG)_n$ to form hairpins. The structures found in the simulations are the result of the generic tendency of C and G to form WC base pairs. Due to the repeat nature of the sequence, the RNA maintains an ensemble of hairpin-like structures by sliding one end over another without paying significant energetic penalty.

The formation of helical structures is also reflected in the bond-bond orientational correlation function,

$$\langle \cos\theta(s) \rangle = \left\langle \vec{b_i}.\vec{b_{i+s}} \right\rangle / l_b^2, \tag{13}$$

where $\vec{b_i}$ is the unit vector of bond $i$, with the bond length $l_b$. $\cos\theta(s)$ shows periodicity at small $s = |i-j|$, where $i$ and $j$ are the indices of the nucleotides in the $(CAG)_n$ sequence (Fig. S3**c**). At a short length scale (up to 5 to 6 nucleotides), the structure of CAG is roughly rigid, with $R(s)$ scaling almost linearly with $s$ (inset of Fig. S3**c**). At larger separations, ($s \geq 6$), $R(s) \propto s^{0.56}$, suggesting that the RNA is flexible, and hence could fold upon itself to generate hairpin-like structures. At a separation corresponding to the end-to-end distance, there is an abrupt decrease in $R(s)$ as a function of $s$ because the 5' and 3' ends are close.

Similar results are also observed for $(CUG)_n$ (Fig. S4). Our simulations show that the hairpin-like structures in $(CXG)_n$ (X = A or U) are a consequence of multiple canonical Watson–Crick base pair formation.

##### Non-canonical base pairing has a minimal effect on the conformations of the repeat monomers

In our model, only canonical Watson–Crick base pairs (bps) are allowed, *i.e.* G only pairs with C, and A only pairs with U. It is known that RNA bases do form a wide variety of other pairings, which may be

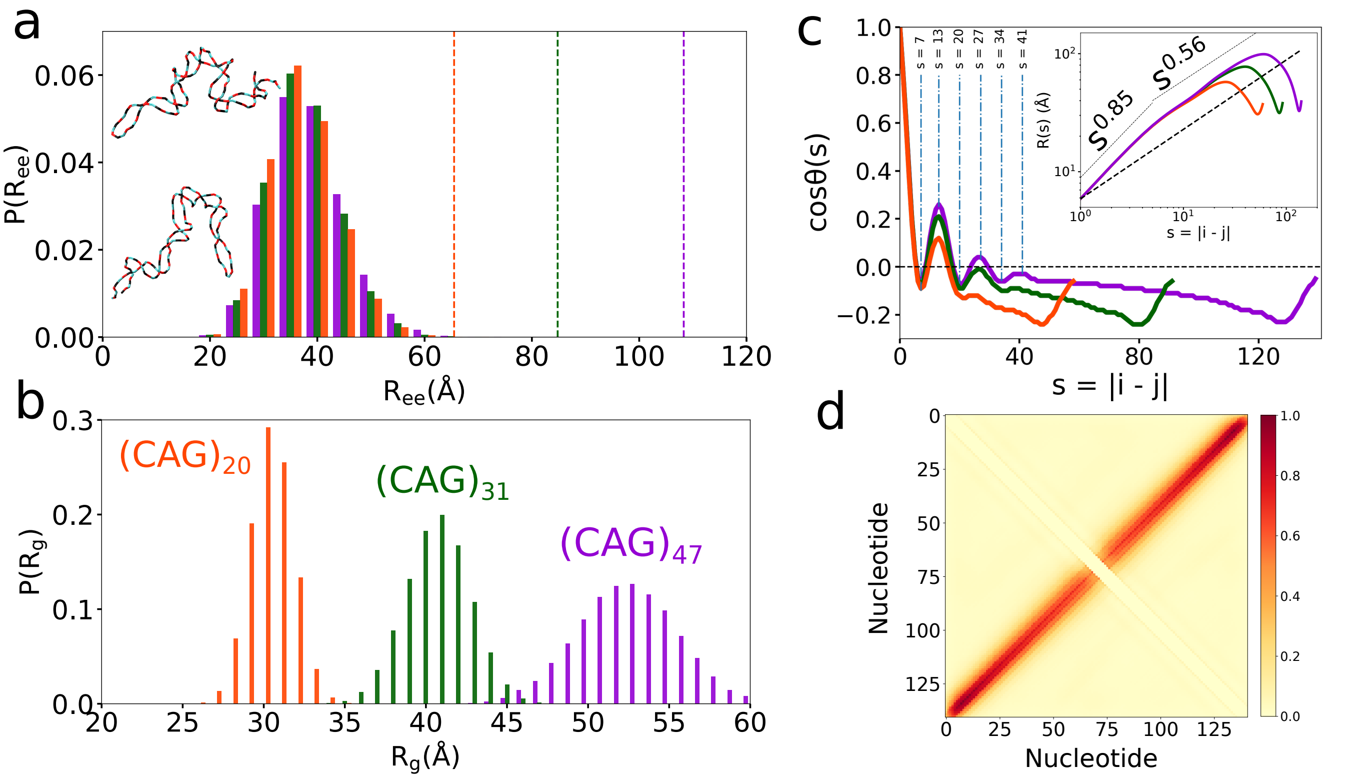


**Figure S3.** Structures of isolated $(\mathrm{CAG})_\mathrm{n}$ monomers. **a**, Distribution of the end-to-end distance $R_{ee}$ and **b**, radius of gyration $R_g$ of $(\mathrm{CAG})_\mathrm{n}$ with n=47, 31 and 20. The vertical dash lines in **a** indicate mean values for self-avoiding random walk chains with the same $n$. Snapshots are for $(\mathrm{CAG})_{47}$. Cytosine is in cyan, adenine is in red and guanine is in black. **c**, Bond-bond orientational correlation function $\cos\theta(s)$ as a function of the sequence distance $s$. The periodicity, as indicated by the vertical lines, is unmistakable. The inset shows average inter-nucleotide distances $R(s)$ vs. $s$. The dashed line shows $R(s)$ for a self-avoiding polymer ($R(s) \propto s^{0.588}$). At large $s$, there is an an abrupt drop in $R(s)$ because the two ends strongly interact with each other, thus bringing them to proximity. **d**, Contact map for $(\mathrm{CAG})_{47}$ shows that the majority of interactions occur along the anti-diagonal, indicating the formation of hairpin structures.

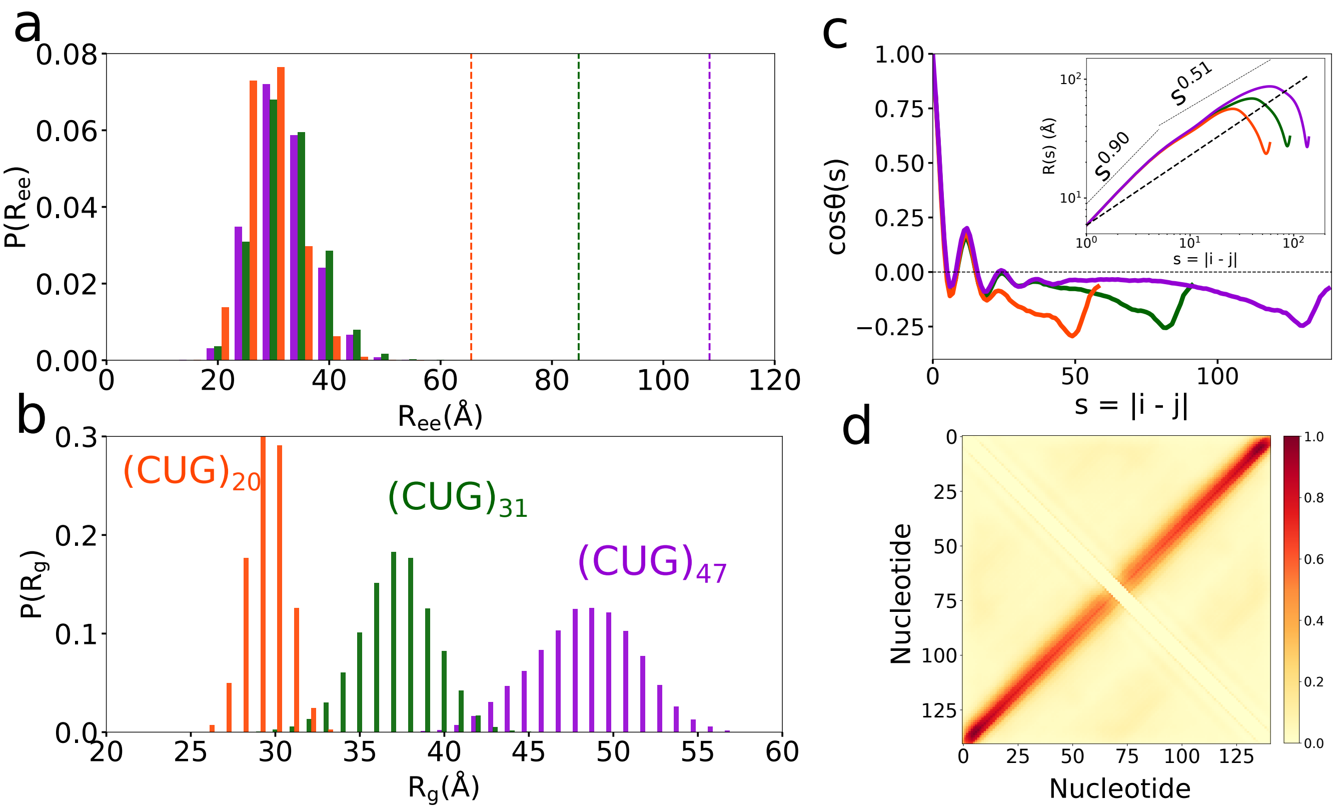


**Figure S4.** Same as Fig. S3, but for $(CUG)_n$ monomers.

classified as non-canonical base pairs (bps).[17] In principle, it is possible to include non-canonical base pairs within our framework. However, by analyzing the RNA structures in the PDB, it is found that the frequency of non-canonical base pair is small compared to WC bps; 70-80% bps in the PDB are WC, and the remaining fraction forms non-canonical bps.[18,19]. Furthermore, while there is only one potential WC bp between two bases (cis-WC-WC, using the Leontis–Westhof notation), there are 11 additional ways to form non-canonical bps by orienting different edges of the base (WC, Hoogsteen or sugar). Therefore, one could infer that the non-canonical bps are much less stable compared to the WC bps, which is the basis for our assumption that WC bp is the dominant force in driving phase separation of the RNA repeats. In addition, the difference between these types of bps mostly arises from the alternative conformations of the base, sugar and phosphate groups within the same nucleotide. Since the SIS model represents a nucleotide by a single bead, any distinction within the nucleotide resolution cannot be modeled explicitly. Therefore, we believe that in order to quantitatively account for the effects of non-canonical base pair formation, one needs the three-interaction-site (TIS) model in which a nucleotide is represented by three beads for the base, sugar and phosphate groups.[7,20–22] The drawback is that a higher resolution naturally raises the computational demand, and making it difficult to simulate multi-chains to probe LLPS using the currently available computing resources.

Having given the justification for neglecting non-canonical bps, here we explored revised the SIS model that includes non-canonical bps between any two bases that cannot form WC bp. The base pairing energy function in the revised model is the same as in the SIS model (Eq. 3), except we include all other possible combinations of bp with a smaller $U^o_{bp} = -1.67$ kcal/mol (which is a third of the value used for a GC bp). When used the revised model to the CAG repeats, this means that A could form non-canonical bps with C, A or G. Even though the modification is a minor addition, it takes much longer

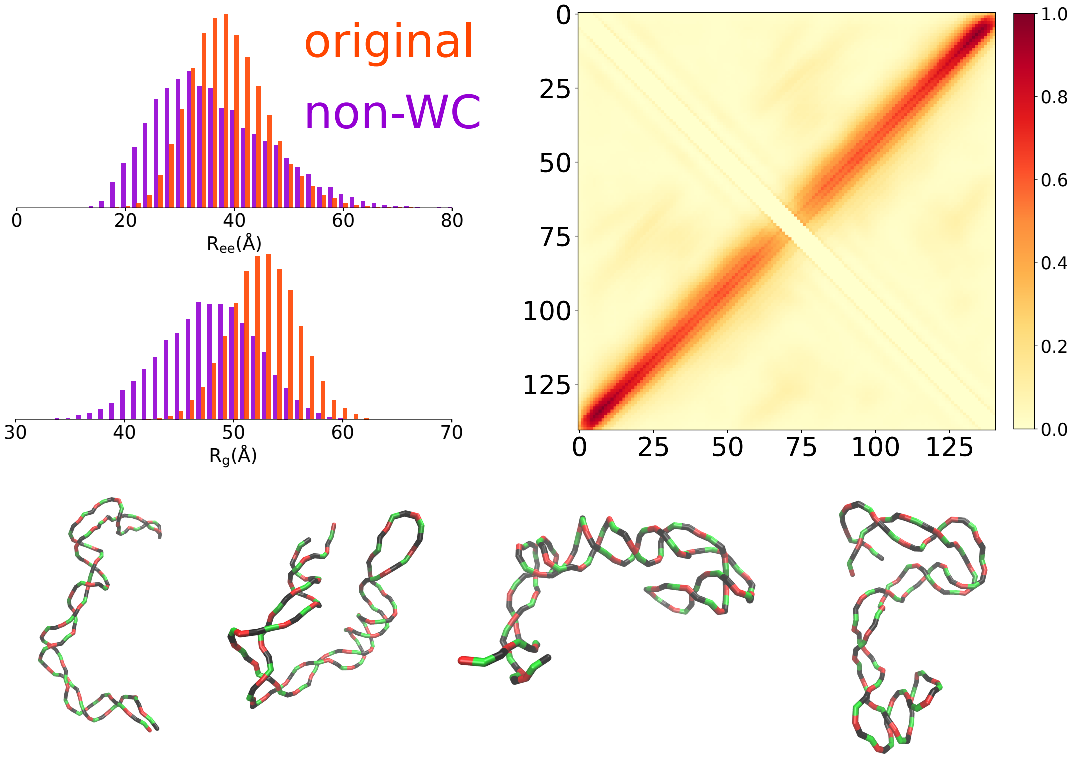


**Figure S5.** Simulations for an isolated $(CAG)_{47}$ monomer where non-canonical base pairs (bps) are allowed (purple), compared with the original model where there are only WC bps (orange). Shown on the left are histograms of the end-to-end distance $R_{ee}$ (top) and radius of gyration $R_g$ (bottom). An intramolecular contact map is shown on the right. Some representative snapshots from the simulations are shown at the bottom.

time to perform simulations compared to the SIS model. Thus, in this preliminary calculation, we report results for the monomer $(CAG)_n$, shown in Fig. S5. The chain is somewhat more compact ($R_{ee}$ and $R_g$ are smaller compared to the SIS model) due to the ability of A nucleotides to form non-canonical bps. However, the conformations are globally hairpin-like, which is exactly the same as in the model that neglects non-canonical bp formation. Therefore, introduction of the non-canonical bp formation in the phase separation simulations could alter the kinetics of hairpin unwinding and the timescale of conversion from intra- to intermolecular bps. However, we anticipate that such modifications would not significantly change the major findings in our work.

#### Effect of cooling rate on droplet formation

An annealing protocol was employed in the *in vitro* phase separation experiments on the CAG repeats.[4] RNA molecules were denatured at 95°C for 3 minutes and cooled at the rate of 1–4°C/min to 37°C. In our simulations, the initial conformations of the RNA molecules are significantly more expanded than in the experiments in order to eliminate biases in the structures that the RNA molecules adopt initially. The expanded initial condition used in the simulations is a mimic of the experimental protocol in which the RNAs are denatured at an elevated temperature for a short duration. Subsequently, the temperature is lowered to the desired value. In order to ensure that our findings are not affected by the initial conditions, we also performed simulations in which we slowly decreased the temperature, as in the experiments. We

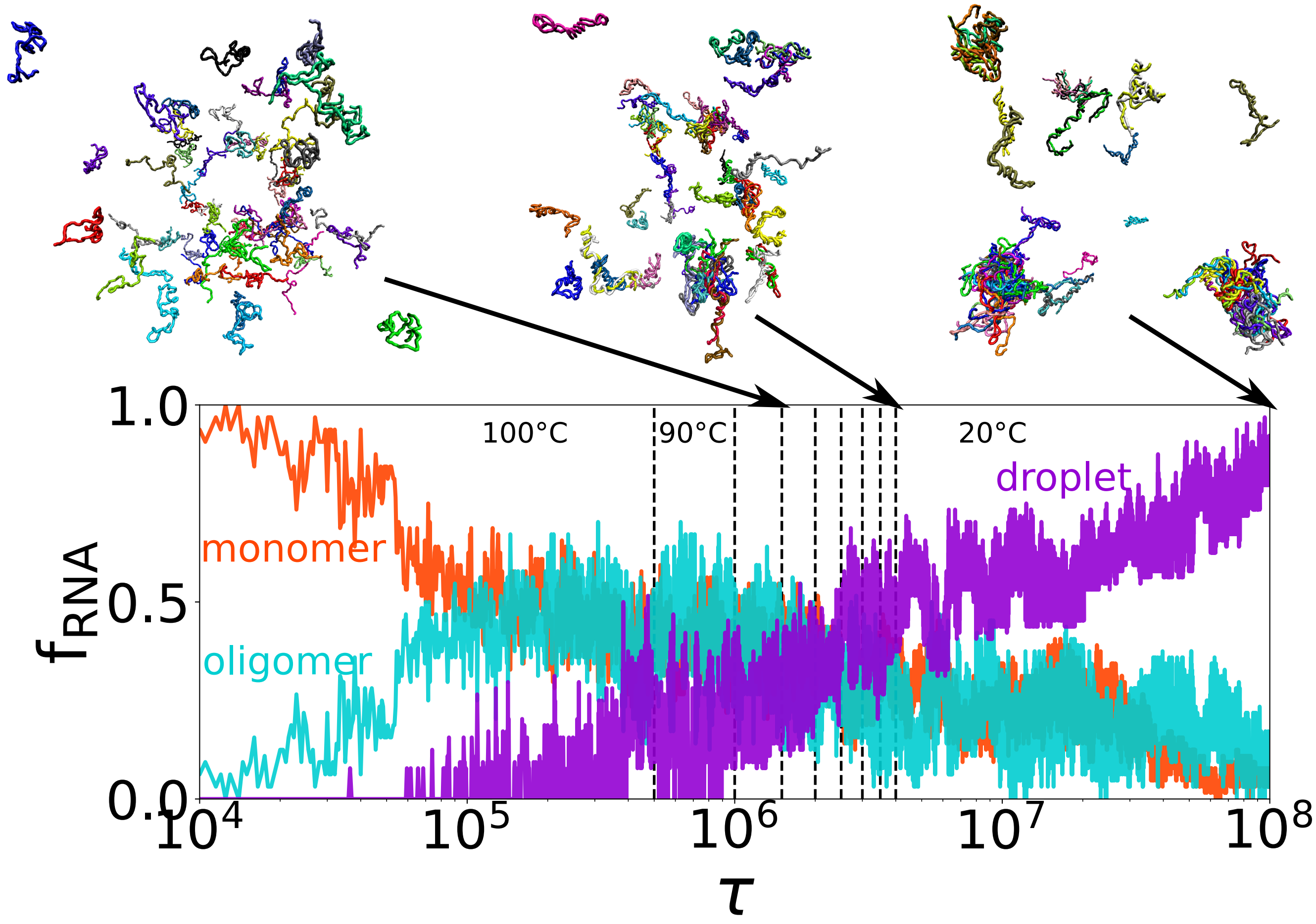


**Figure S6.** Condensate formation does not depend on the cooling rate. Fraction of chains in droplets or existing as oligomers/monomers. The vertical dashed lines indicate when the temperature is lowered (from 100°C to 20°C). Snapshots from left to right correspond to, respectively, the end of 80°C, the end of the cooling period and the final state of the simulation.

first ran simulations at 100°C, which is high enough such that all the RNA molecules are unfolded. Then, after every $\delta t$, we lowered the temperature by 10°C until the final temperature reached 20°C. We could explore the cooling rate by varying $\delta t$. We hasten to add that limitations in the simulation times prevent us from reaching the cooling rates achieved in experiments. Fig. S6 shows that the overall picture of phase separation is not qualitatively altered, regardless of the procedures followed in initiating the simulations. It indicates that our simulations are robust with respect to the initial conditions and simulation protocols, and that the phase separation of the repeat RNAs occurs spontaneously, just as in experiments. It should be noted that experimental studies from other groups also showed that RNA molecules alone undergo phase separation without using the aforementioned annealing method,[23,24] or using a rapid cooling rate as we did here[25]. Thus, it appears that initial conditions do not affect the mechansism of LLPS in RNA repeat sequences.

##### Droplet stability at low salt concentrations

To probe the droplet formation in low salt conditions, we included electrostatic effects. In addition to the bond, angle, excluded volume and hydrogen bond terms as in Eq. 1, each nucleotide now carries a

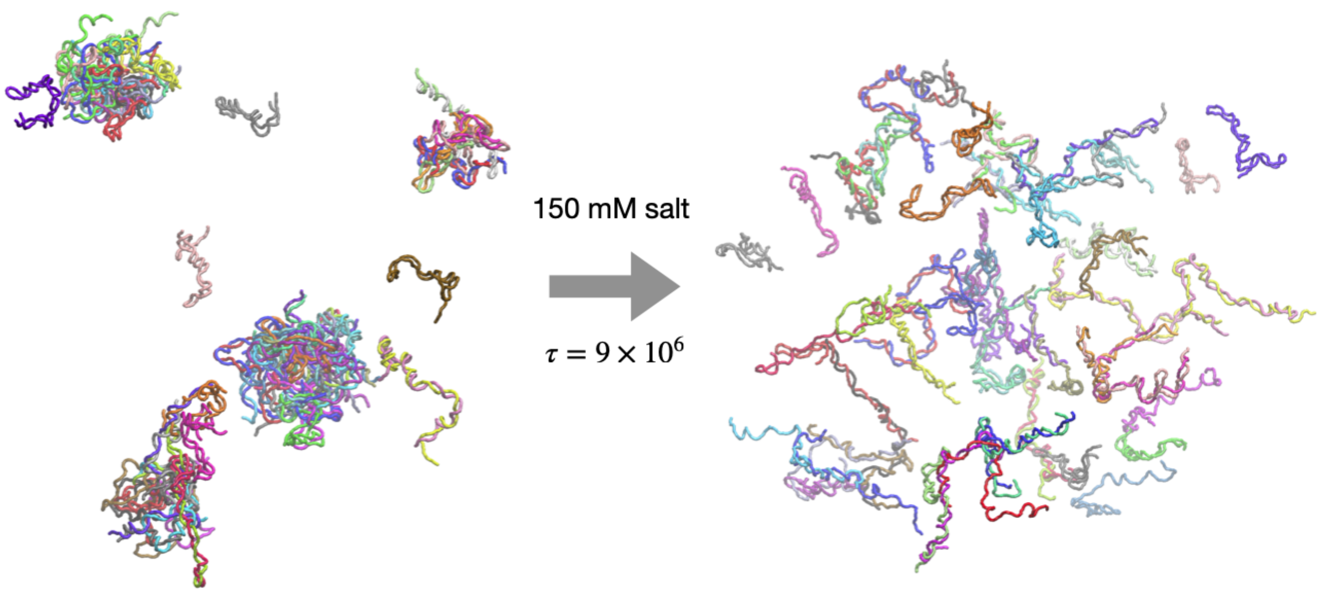


**Figure S7.** Dissociation of RNA droplets at 150 mM NaCl. The simulations were started from the final configuration obtained in the droplet simulation of 200 $\mu$M of $(CAG)_{47}$ (left). The repulsive electrostatic interactions between RNA nucleotides (due to the incomplete neutralization of phosphate charges) lead to the disassembly of the droplets, leaving only monomers and small oligomers (right).

charge of $-q$ $(q < 1)$ to account for counterion condensation. The electrostatic repulsion between them is modeled using the Debye–Hückel theory, accounting for the screening effect of the buffer solution. This approach has been successfully used to calculate accurately the thermodynamics of RNA folding.[7,20] To test if the phase separation occurs at low salt conditions, we performed simulations using a model that includes ion effects. The initial conditions correspond to a preformed condensate. The simulations show that the RNA droplets start dissociating into smaller droplets to form oligomers and monomers. Thus, in the presence of monovalent ions alone, the droplet is unstable, which recapitulate qualitatively the experimental findings.

Of course, these simulations using the DH theory only accounts for the effect of monovalent ions ($Na^+$, $K^+$ ...). This approach could be extended to include divalent cations ($Mg^{2+}$ ...) explicitly, while keeping monovalent ions at the continuum level, as we have shown previously for the RNA folding problem.[7] Although simulations using such a model would be computationally demanding, they would provide the complete phase diagram. The SIS model is sufficiently general to accommodate ion explicitly, as shown elsewhere.[7,20]

#### Scrambled sequences are less likely to undergo LLPS

We then tried a different sequence to test the transferability. We shuffled the sequence of $(CAG)_{47}$ to maintain the GC content and chose the sequence (out of an astronomically large number of sequences) C CGGGAAGAGACCGCAACAGAAGCAGCCGCGAGCGCGACAGCGACGAGCACGCGGCACA GACAGCAAGAGAAGGGAAGACAAAGAGCCGACGGAAGCCACGCAGAGCAAAAGGACGC CGGCGGAACGACAAGGAAAGAGAG. To the best of our knowledge, Jain and Vale did not report the order of nucleotides in their scrambled sequence, which forced us to create one for computational purposes. We then repeated the simulations at different bulk concentrations 200, 100, 50 and 20 μM. We observed that this particular scrambled sequence undergoes phase separation, which seems to be

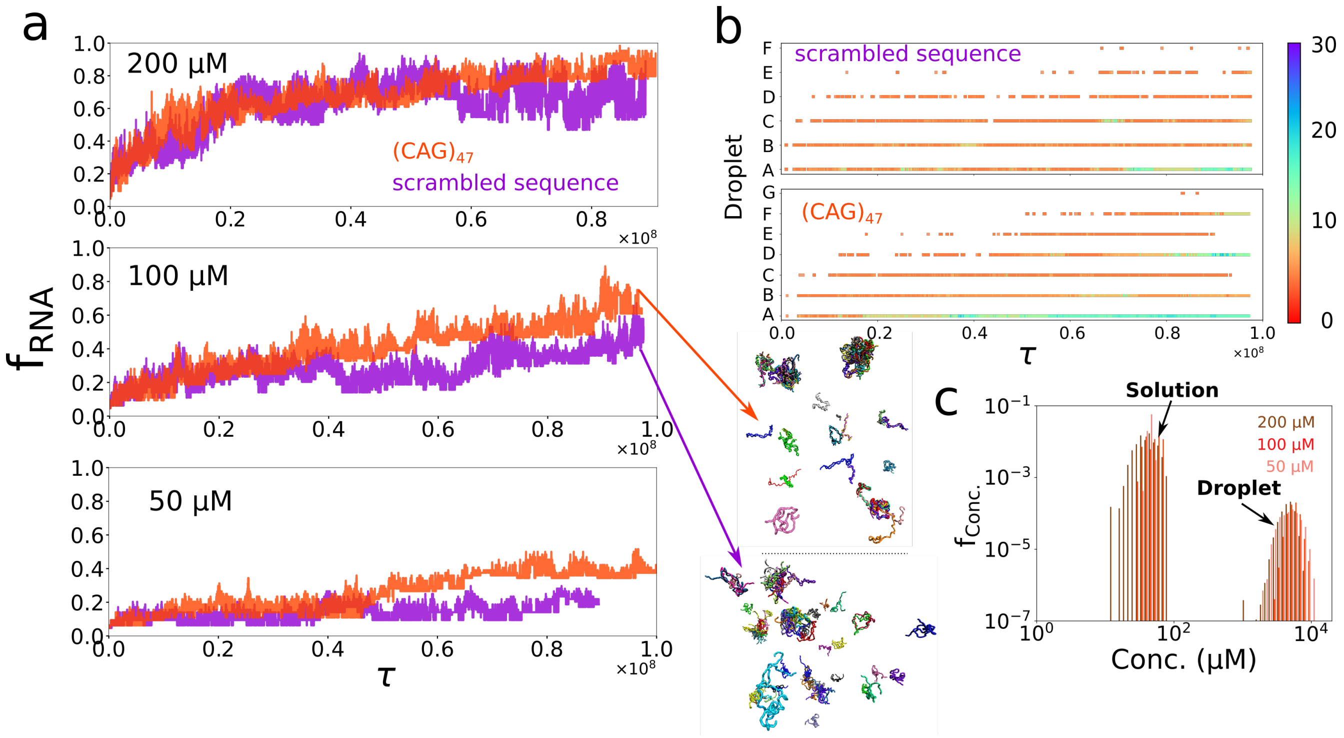


**Figure S8.** Simulations for the scrambled sequence. **a**, Comparison of fraction of RNA chains inside the droplets for $(CAG)_{47}$ and the scrambled sequence at three different concentrations. Snapshots near the end of the simulations for the two sequences are shown. **b**, Droplet size evolution for the scrambled sequence (top) vs. $(CAG)_{47}$ (bottom). Each horizontal line corresponds to a specific droplet in the system. The size is denoted by the color (color scale is on the right). **c**, Concentrations of the two phases for the scrambled sequence.

in apparent contradiction with the finding in the Jain–Vale experiment. However, in our simulations, the fraction of chains that are in the droplet state for the scrambled sequence is lower than in $(CAG)_{47}$ (Fig. S8**a**). As a result, the concentration of the diluted solution in the case of the scrambled sequence is larger (45 μM, compared to 25 μM for $(CAG)_{47}$). We emphasize that the concentration of the diluted phase is the saturated concentration $C_{\text{sat}}$, above which phase separation occurs. The observation of $C_{\text{sat}}^{\text{scrambled}} > C_{\text{sat}}^{(\text{CAG})_{47}}$ indicates that the propensity to undergo phase separation of the scrambled sequence is lower than in $(CAG)_{47}$. On the other hand, the concentration inside the droplets of the scrambled sequence is smaller than droplets composed of $(CAG)_{47}$ (5 vs. 10 mM), suggesting less compaction of the scrambled sequence droplets. This decreased compaction is likely due to the random nature of the sequence, leading to a decreased probability to find similarly patterning neighboring chains to form base pairs. We also observed that the droplet sizes in the case of the scrambled sequence are smaller than $(CAG)_{47}$, and the system tends to form many smaller droplets instead of a few relatively large droplets.

Taken together, the new simulations suggest that the propensity to undergo LLPS of the scrambled sequence is less than the repeat sequence, which may be in qualitative agreement with the experiment. It is possible that a much higher concentration of the scrambled sequence is needed to induce phase separation. Even if the concentration is higher than $C_{\text{sat}}$, it is conceivable that due to the smaller droplet sizes in the scrambled sequence compared to the repeat sequence, it is below the critical threshold detectable in the experiment (or one would need to wait longer for the droplets to grow to detectable sizes).

#### Local behavior of RNA chains growing from monomer to oligomer to droplet

A major advantage of simulations is that the microscopic behavior of the RNA chains can be visualized and investigated, if the simulations are validated against experiments. In particular, it is interesting to probe the sequences of events, which are not currently accessible in experiments, that occur as the monomer becomes part of the condensate. One difficulty is that the fate of each chain could be different, which implies that there may not be typical or most probable changes in the dynamics as the monomer becomes part of the droplet. With this caveat, we performed several analyses focusing on the local behavior of chains.

Figure S9 illustrates examples of local dynamics while multiple chains form a single droplet. In the series of snapshots, we show a process of coalescence by eleven RNA chains that eventually form an 11-mer in the middle of the simulation at 200 $\mu$M $(CAG)_{47}$. We observed that chains often form small oligomers first, then they coalesce to form larger oligomers, leading eventually to a droplet. Oligomers can also grow by merging with monomers and dimers. In the sample trajectory, we show that a trimer and a tetramer coalesce into a heptamer, followed by integrating other dimers to be an 11-mer at last. A more detailed description of the event is in the caption of Figure S9.

#### Reptation does not occur for monomers

Fig. S10 shows MSD for nucleotides in an RNA monomer in the diluted phase. The MSD scales as $\tau^{0.78}$, which is significantly larger than $\tau^{1/4}$ expected for reptation. This proves that reptation dynamics only occur inside the droplets.

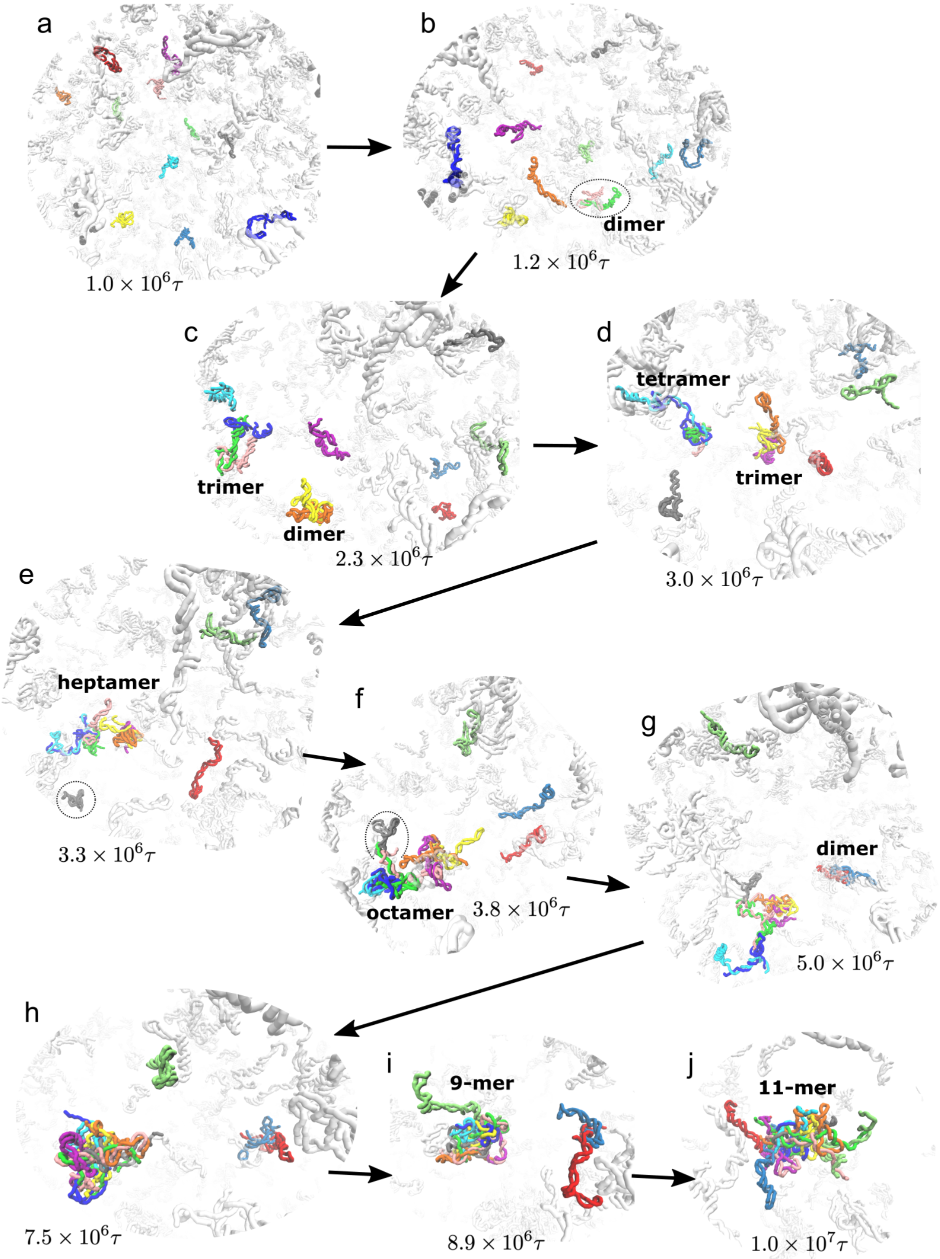


**Figure S9.** Sequence of events in early droplet formation extracted from the simulation of $(CAG)_{47}$. Eleven RNA chains were chosen and colored to see how individual chains form oligomers and grow to a single droplet. All other chains are in grey for clarity. Each panel has a label indicating the simulation time. (a) At the earliest times, all the eleven chains are monomers with no interactions between them. (b) Two chains merge to form a dimer. (c) The dimer captures another chain and becomes a trimer. There is another dimer that is formed around the same time. (d) The two oligomers further grow to a tetramer and trimer, respectively, by interacting with another chain. (e) The tetramer and trimer coalesce into a heptamer. There are still four other chains in the monomer form. (f) One of the remaining monomers joins the oligomer making it an octamer. (g) Two of the remaining monomers form a dimer. (h) It takes some time to the next event ($\sim 4 \times 10^6 \tau$ from (g) to (h)). (i) The octamer eventually captures the last monomer and becomes a nonamer. (j) The nonamer and the dimer finally coalesce into an 11-mer. The sequence of events is complicated, and is different for different chains.

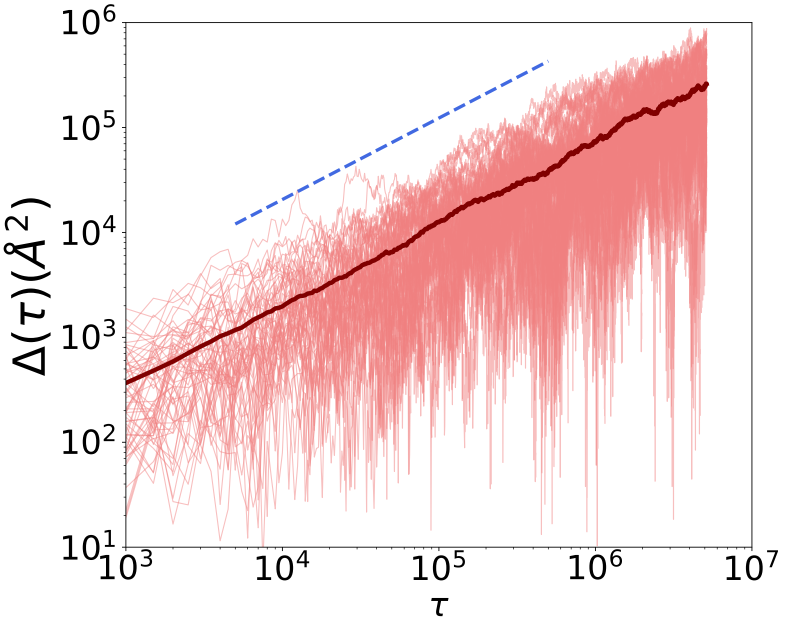


**Figure S10.** Mean square displacement (MSD) of nucleotides in monomers in the diluted phase. The average values are plotted as the solid line. The MSD scales as $\tau^{0.78}$ (blue) at all time scale, showing no sign of reptation dynamics.

#### Supplementary Figures

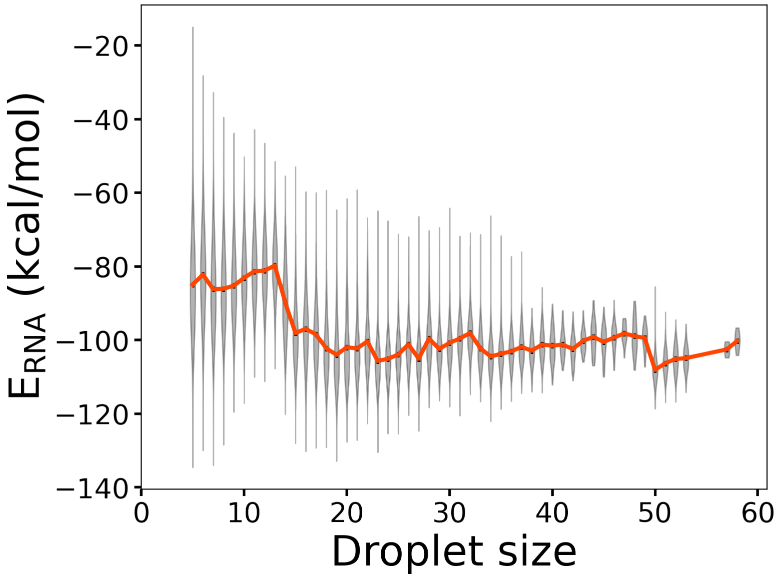


**Figure S11.** Energy per chain vs droplet size. As the size of the condensate increases, the energy per chain decreases, which indeed is the reason for phase separation from a thermodynamic perspective.

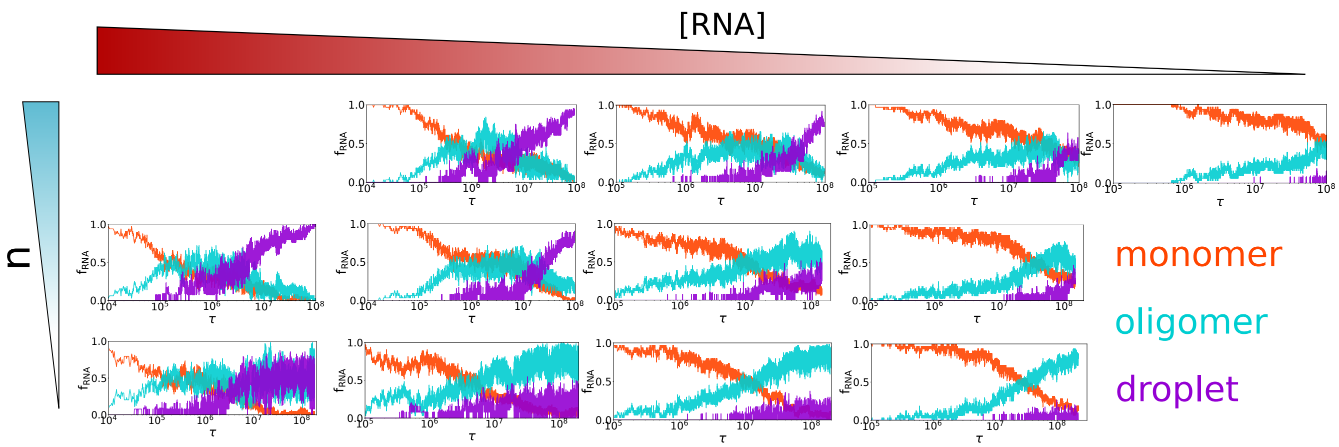


**Figure S12.** Fraction of monomers, oligomers and droplets as a function of (CAG) repeat length and concentration.

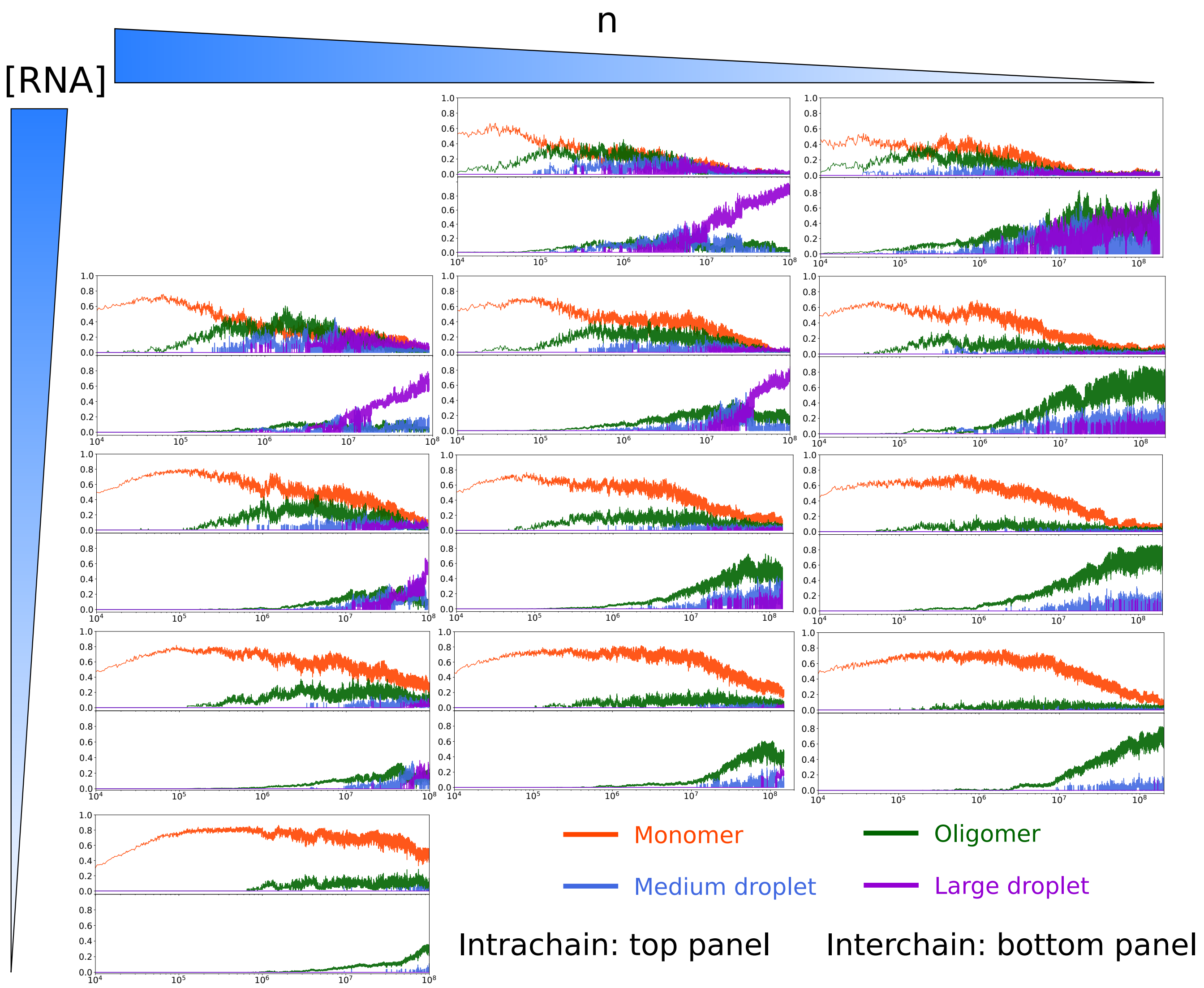


**Figure S13.** Fraction of (both intra- and intermolecular) hydrogen bonds as a function of time. The conversion of intra- to intermolecular interactions is the dominant factor driving the phase separation.

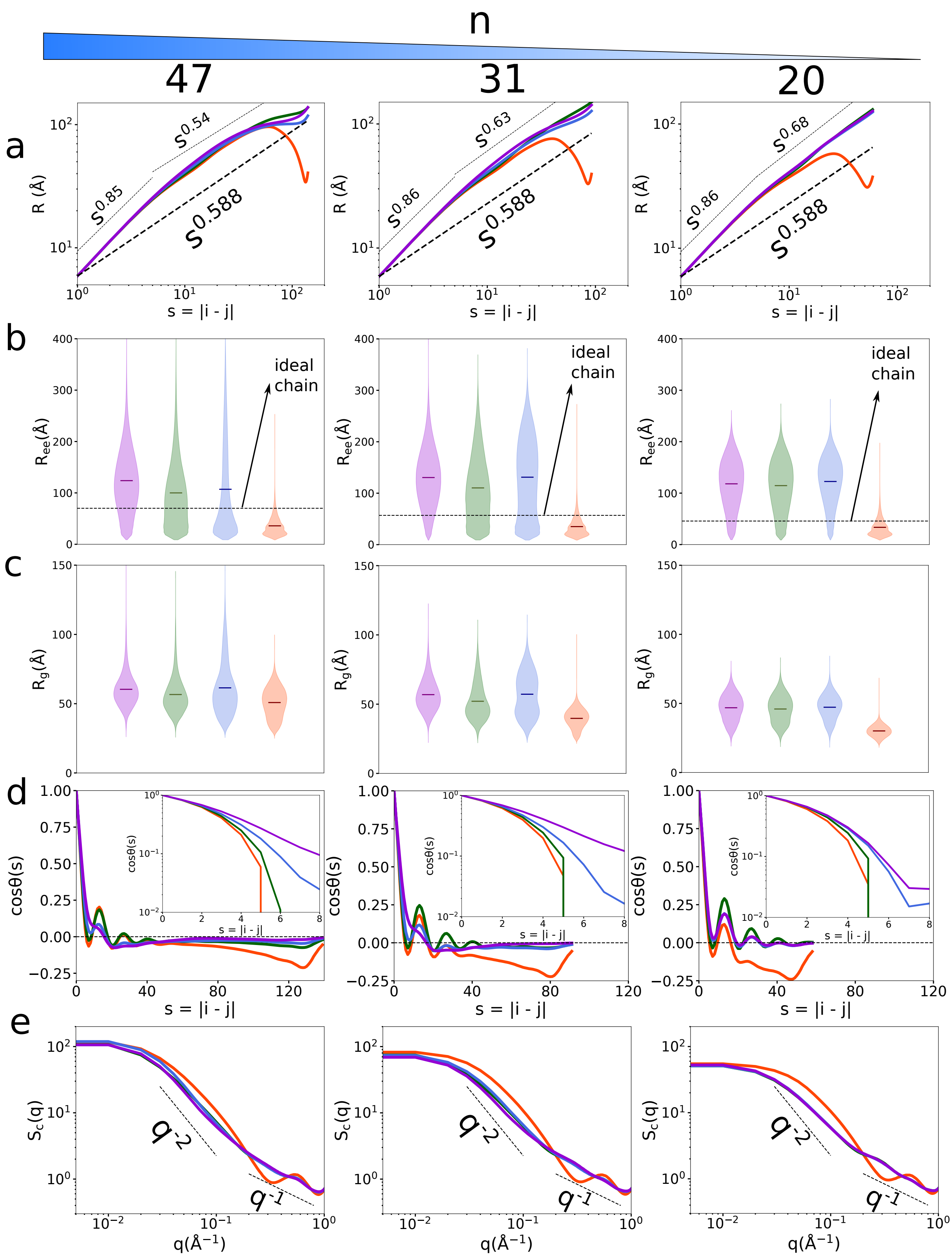


**Figure S14.** RNA undergoes significant conformational changes upon entering droplets.

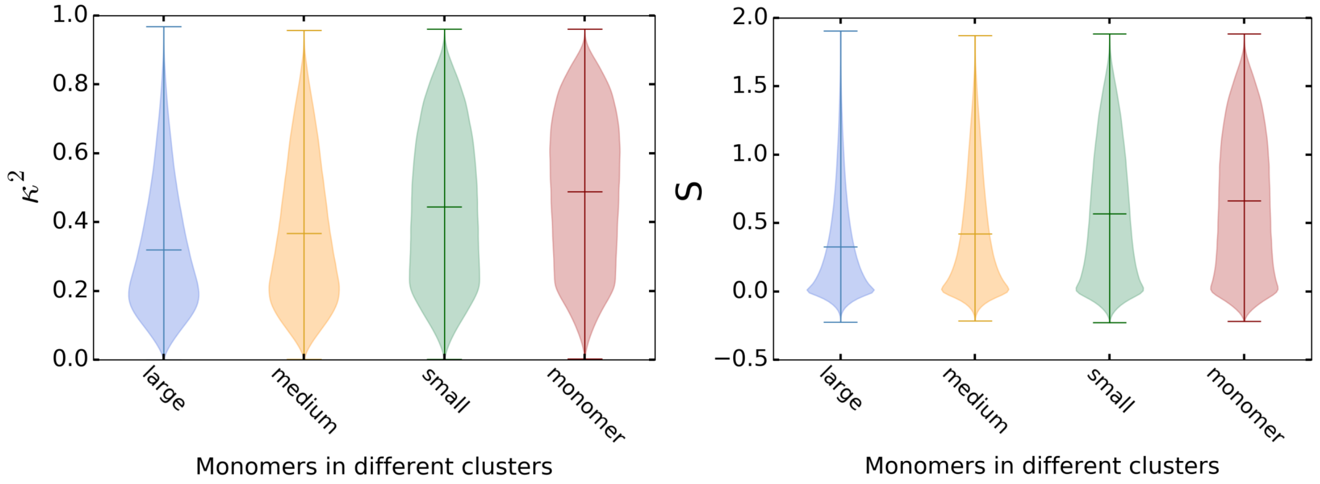


**Figure S15.** Comparison of shape parameters of monomeric RNAs in solution and in droplets.

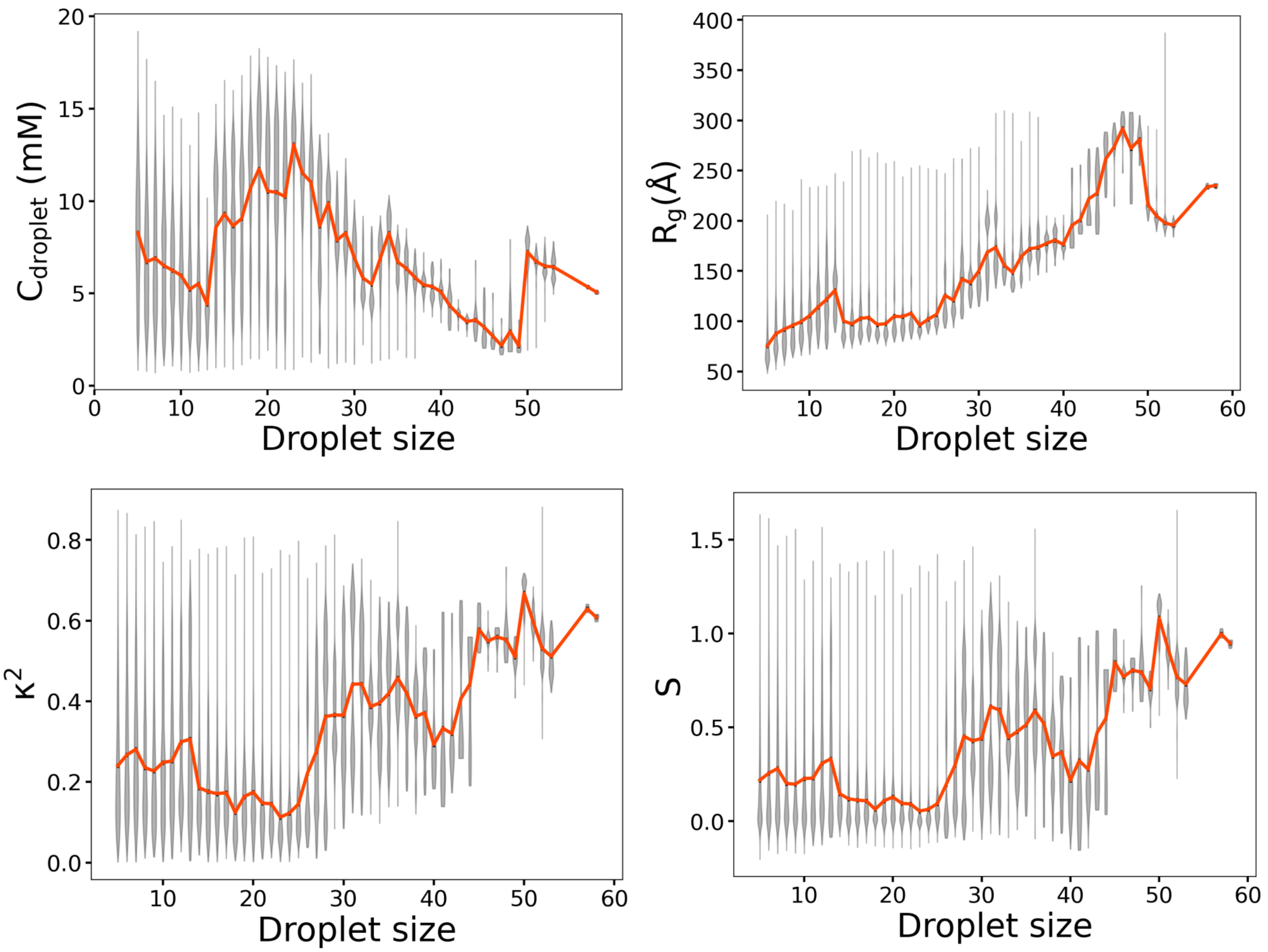


**Figure S16.** Characterization of droplet properties as a function of droplet size.

#### Supplementary Movies

**Movie S1.** Dynamics of phase separation in $(CAG)_{47}$ from monomers to condensates. The monomer RNA concentration is 200 $\mu$M. At the initial time ($\tau = 0$), the chains exist as monomers. At intermediate times, oligomers form, which subsequently fuse together resulting in large droplets as time progresses. See Figures 1a, 3a, and 5 for the corresponding trajectories. Note that some of the individual RNA chains may look less helical shape due to trajectory-smoothing needed for easier visualization. The movie shows both fusion as well as fission of the droplets.

**Movie S2.** Growth and fusion of condensates. The movie shows two small clusters of RNA chains (green and magenta) fuse to form a droplet. In the process, the cluster in magenta is once partially dissolved but eventually coalesce with the other cluster (green).

**Movie S3.** Jamming dynamics of two labelled RNA chains inside a droplet. The movie shows the late-stage dynamics in a simulation trajectory at 200 $\mu$M $(CAG)_{47}$ after the formation of a large droplet. For clarity, focus is on two RNA chains shown in yellow and red colors, whereas all other chains in the same droplet are shown as transparent blue chains. The motions of these RNA chains are highly restricted, resulting in reptation-like dynamics (Figure 7).
